# Nitrogen use efficiency in pigs is associated with transcriptomic signatures related to amino acid metabolism, immune activity, and nutrient partitioning

**DOI:** 10.64898/2026.06.26.733976

**Authors:** Bastien Monney, Esther Oluwada Ewaoluwagbemiga, Claudia Kasper

**Affiliations:** Animal GenoPhenomics group, Agroscope, Posieux, Switzerland

**Keywords:** differential gene expression, pathway enrichment, protein turnover, mitochondrial metabolism, tissue-specific regulation

## Abstract

Dietary protein restriction challenges the allocation of amino acids to growth and other physiological functions and therefore requires coordinated metabolic adaptation. Domestic pigs provide an informative system in which to study such responses, because nitrogen retention directly affects lean growth and can be quantified accurately under controlled feeding and housing conditions. Under reduced-protein diets, pigs differ in how effectively they retain nitrogen, and this variation has a genetic basis, making them well suited to investigate the molecular regulation of nitrogen use efficiency (NUE). Here, we characterise differential gene expression and enriched pathways in liver and skeletal muscle of more than 80 pigs with two divergent NUE phenotypes (high and low) maintained under the same protein-reduced, *ad libitum* dietary conditions. The two NUE phenotypes were clearly distinct at the transcriptomic level, with 177 differentially expressed genes in the liver and 133 in the muscle. In the liver, differential expression and enrichment analyses indicate reduced amino acid catabolism, lower inflammatory and detoxification activity, and a metabolic state that favours lipid processing and insulin-related regulation over the use of amino acids as energy sources. In skeletal muscle, they point to reduced lipid uptake, lower reliance on amino acid oxidation, and a greater emphasis on protein synthesis, translational regulation, mitochondrial energy metabolism, and growth-related processes. These gene-level patterns were supported and extended by pathway and gene-set enrichment analyses. Together, the results suggest that high and low-NUE pigs differ through coordinated, tissue-specific molecular adaptations. Overall, variation in NUE appears to reflect coordinated, tissue-specific differences in how nutrients are allocated between energy use, storage, and lean tissue growth.

## Introduction

Dietary protein availability is a fundamental determinant of mammalian growth, tissue maintenance, and metabolic homeostasis. Amino acids are required not only for the accretion and maintenance of body proteins, particularly in skeletal muscle, but also the synthesis of a wide range of biologically active molecules, including nucleotides, hormones, and components of the immune system such as antibodies, and neurotransmitters. When protein supply is limited, their allocation across these competing functions becomes a central physiological challenge. Given their role in the tight regulation of protein synthesis, amino acid oxidation, and nitrogen excretion, a limited protein supply can trigger the conservation of amino acids through coordinated cellular responses, such as suppression of anabolic pathways such as mTORC1 signalling and activation of autophagy and the integrated stress response (Bröer and Bröer, 2017). At the organism level, the maintenance of amino acid homeostasis depends particularly on the coordinated functions of multiple tissues, particularly the liver and skeletal muscle: the liver is a central site of amino acid catabolism, ureagenesis, and systemic amino acid regulation (Wester et al., 2015), whereas skeletal muscle constitutes the largest protein reservoir in the body and a major site of protein turnover and accretion (Wolfe, 2006). Accordingly, a limited protein supply requires coordinated adjustments in nutrient sensing, amino acid metabolism, and tissue-level resource allocation to maintain metabolic function and support growth.

Domestic pigs provide a particularly informative system for the study of the biological consequences of variations in dietary protein use, because the efficiency with which nitrogen is retained in body tissues directly affects both lean growth, which is central to pork production, and nitrogen excretion in manure, a major source of reactive nitrogen emissions with adverse environmental effects (Kasper, 2024). In addition, pigs offer a tractable system for studying individual variations in nitrogen retention, because controlled feeding and housing conditions, as well as established phenotyping protocols allow this trait to be quantified with a level of accuracy that is difficult to achieve in most other mammals, including humans. This efficiency is commonly described as nitrogen use efficiency (NUE), that is, the proportion of dietary nitrogen that is converted into body protein rather than lost into the environment through excretion. Improving NUE is therefore relevant not only for reducing the environmental footprint of pig production, but also for understanding how animals sustain growth when the protein supply is limited. Reducing dietary crude protein is a widely used strategy to decrease nitrogen excretion in pig production, particularly when the amino acid supply is balanced according to the animals’ requirements (Liu et al., 2024; Pomar et al., 2021). However, animals differ in how effectively they retain nitrogen in the body and maintain growth under such conditions (Kasper et al., 2020; Ruiz-Ascacibar et al., 2019), and previous work has shown that this variation in NUE has a genetic basis (Ewaoluwagbemiga et al., 2023).

Transcriptomic studies of pigs that differ in terms of broader feed efficiency traits provide further evidence that efficiency-related phenotypes are associated with coordinated, tissuespecific differences in metabolic and signalling pathways, especially in the liver and skeletal muscle (Horodyska et al., 2019, 2018; Vigors et al., 2019). In particular, these studies have pointed to shifts in how nutrient and metabolic substrates are allocated between storage, mobilisation and energy production, including altered lipid and carbohydrate metabolism, as well as differences in cellular growth and cellular communication, consistent with the view that efficiency reflects a broader metabolic reorganisation rather than changes in isolated pathways alone (Reyer et al., 2017). Similarly, pigs from a divergent selection experiment for feed efficiency exhibited distinct transcriptomic and proteomic profiles in skeletal muscle, with efficient animals generally showing higher expression of genes involved in protein synthesis and lower expression of genes and proteins involved in oxidative metabolism (Vincent et al., 2015).

However, whether similar tissue-specific transcriptional patterns are associated specifically with NUE under protein-limited conditions remains largely unexplored. This is important because efficiency-associated molecular profiles may differ across nutritional environments, indicating potential genotype-diet interactions. Resolving this question is relevant both for understanding biological variations in performance under a limited protein supply and for improving complementary mitigation strategies to reduce nitrogen losses. In practice, such strategies are likely to require the integration of nutritional and genetic approaches, including reduced-protein feeding and selection for improved NUE. This, in turn, requires a clearer understanding of the genes and biological pathways underlying inter-individual differences in nitrogen retention under protein-limited conditions. NUE reflects the integrated action of multiple interdependent physiological processes across tissues and is shaped by both animal-intrinsic and environmental factors; hence it is expected to be a complex, polygenic trait (Kasper, 2024). In line with this, previous genomic analyses have shown that NUE has a polygenic architecture and is influenced by many loci with small effects, making its biological basis difficult to resolve through association mapping alone (Ewaoluwagbemiga et al., 2025). As expected, these analyses did not identify major loci that readily explain variation in NUE, but highlighted several plausible candidate genes and pathways, including nutrient sensing, the urea cycle, and broader metabolic pathways, particularly the IGF1–insulin axis. Tissue-specific transcriptomic data are therefore needed to connect thes diffuse genetic signals to the underlying biological processes and to identify the genes and pathways most relevant to nitrogen retention under protein-limited conditions.

To address this, we analysed the liver and skeletal muscle gene expression of pigs differing in NUE that were maintained under protein-limited conditions with a balanced essential amino acid supply and no caloric restriction, all within the same nutritional environment. Together, these tissues provide complementary insights into the biology of nitrogen retention and growth. By combining gene-level and pathway-level analyses, we aimed to identify transcriptional differences associated with variations in NUE, tissue-specific expression patterns across functionally related genes, and biological processes linked to efficient nutrient allocation under a limited protein supply.

## Materials and methods

### Animals

All pigs used in this study were reared in an experiment to investigate genetic parameters of NUE in pigs (Ewaoluwagbemiga et al., 2023), which also formed the basis for a genome-wide association study (Ewaoluwagbemiga et al., 2025). All experimental procedures were approved by the Swiss Federal Food Safety and Veterinary Office (30714/2018-30-FR) and conducted in accordance with Swiss regulations on animal welfare and animal experimentation. The animals were Large White dam-line pigs reared at the experimental farm at Agroscope Posieux under controlled feeding and housing conditions with individual feed intake recording. From the start of the experiment until slaughter, pigs received protein-reduced grower and finisher diets containing 80% of the crude protein and digestible essential amino acids recommended for pigs in Switzerland (Stoll, 2004). The samples analysed here represent a subset of these pigs selected according to NUE and sex (see section “Choice of samples for RNA-seq” below). Further details on animal management, diet composition, phenotyping, and NUE calculation are provided in the original publications. The dataset used here is available online (Monney et al., 2026).

### Model variables and calculation of nitrogen use efficiency

The models used for the differential gene expression analysis included NUE and routinely collected production variables as covariates. These comprised sex, birth weight, slaughter weight, age at slaughter (in days), and ambient temperature, feed conversion ratio (FCR), feed intake and growth performance (Table 1). These variables were included to account for biological and management-related variation within the study population. This was also relevant because, in the larger cohort from which the present subset was drawn, several production traits, including ADG, ADFI, and FCR, were correlated with NUE at both the genetic and phenotypic level (Ewaoluwagbemiga et al., 2023). The feed intake was continuously recorded via automated feeder stations, which logged each pig’s visit. Total feed intake was calculated from these records, and average daily feed intake (ADFI) was derived by dividing the total intake by the number of days on feed. The average daily gain (ADG) was calculated as

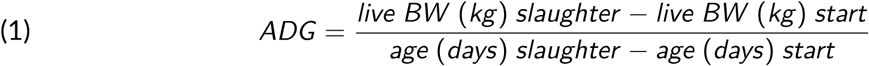

**Table 1.** Variables recorded on pigs with their definition and ontology terms

| Variable | Abbreviation | Measurement unit | Definition | Ontology term |
| --- | --- | --- | --- | --- |
| Nitrogen use efficiency | NUE | - | The amount of nitrogen output for production (i.e., in the carcass) to the amount of nitrogen intake from feed | ATOL_0001583 |
| Average daily gain | ADG | kg/day | Difference in live body weight at the beginning and the end of a defined period divided by the duration of the period in days | ATOL_0000989 |
| Average daily feed intake | ADFI | kg/day | Total amount of feed consumed over a defined period divided by the duration of the period in days | ATOL_0005508 |
| Feed conversion ratio | FCR | - | Ratio between body weight gain and feed intake over a defined period of time | ATOL_0001580 |
| Sex | - | - | Sex of the pig (female or castrated male) | ATOL_0000403 |
| Birth weight | - | kg | Body weight at birth | ATOL_0000093 |
| Slaughter weight | - | kg | Body weight at slaughter | ATOL_0000351 |
| Age at the start of experiment | - | days | Age of the pig in days counted from birth at the start of the experiment (approximately at a body weight of 22 kg) | ATOL_0000088 |
| Age at slaughter | - | days | Age of the pig in days counted from birth at slaughter (approximately at a body weight of 106 kg) | ATOL_0001542 |
| Average ambient temperature | - | °C | Average of the air temperature in the barn of the period starting three days before and after the pig started the experiment | EOL_0000219 |

where live BW (kg) slaughter and age (days) slaughter are the live pre-slaughter body weight in kg and the age in days at slaughter, respectively, and live BW (kg) start and age (days) start are the exact body weight in kg and the age in days at the start of the grower phase, respectively. FCR, defined as. the amount of feed required to produce one kg of live weight, was calculated as

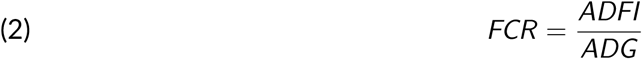

To obtain NUE, the lean tissue content of the left carcass half was determined using dual-energy X-ray absorptiometry (DXA; GE Lunar i-DXA, GE Medical Systems, Glattbrugg, Switzerland) and converted into crude protein content via the equation

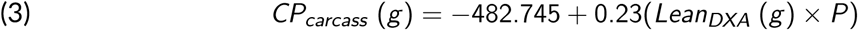

where CPcarcass (g) is the crude protein content of the carcass, LeanDXA (g) is the lean meat content from DXA, and P is the proportion of the left cold carcass (includes the whole head and tail) of the total cold carcass weight (Kasper et al., 2021). Finally, NUE was calculated as

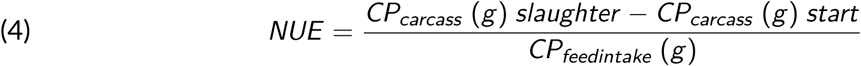

where CP*_carcass_*(g) is the crude protein content of carcass at slaughter, CP*_carcass_* (g) start is the estimated crude protein content of carcass at the start of the experiment, and CP*_feedintake_* (g) is the crude protein intake over the experiment. CP*_carcass_* (g) start was estimated from a sample of 12 female and 12 castrated male piglets slaughtered at approximately 20 kg in a previous experiment. To account for environmental influences, the average ambient temperature in the barn at the start of the experiment (calculated as the mean over three days before and after each pig entered the trial) was included in the model. This was based on previous evidence showing that elevated ambient temperatures can reduce NUE by limiting protein retention in growing pigs (Berghaus, 2022).

### Slaughter and tissue collection

Upon reaching the target slaughter weight, the pigs were scheduled for slaughter the following week at the Agroscope abattoir. They were slaughtered after a feed withdrawal period of 16 h. They were stunned for 180 s with a CO_2_ (87% CO_2_) stunner (Samson C1 L 803; MPS Group, Holbaek, Denmark) and immediately exsanguinated. After scalding at 62°C for 3 min and removal of the hooves, the carcasses were eviscerated. Approximately 15-20 min after exsanguination), a 1 g sample was taken from both the longissimus thoracis muscle and the right lobe of the liver. All samples were immediately snap-frozen in liquid nitrogen. Samples were stored at -80°C until RNA isolation.

### Choice of samples for RNA-seq

The RNA-seq subset was selected to maximise the contrast between animals with divergent NUE phenotypes, thereby providing an informative basis for discovery-oriented transcriptomic analysis. As NUE is a continuous trait, pigs were selected from the high and low ends of the NUE distribution. The final selection of livers and skeletal muscles was performed separately, depending on tissue availability, RNA quality and achieving a balanced representation of males and females. Due to logistical constraints during the pandemic, particularly staff shortages at the time of slaughter, liver and/or muscle tissue could not be collected from some animals. To define the high- and low-NUE groups, the pigs from the full population described above were ranked by NUE, which averaged 0.393 ± 0.030 (mean and standard deviation [SD]). Animals with high NUE were defined as those exceeding the mean plus one SD (> 0.423), and those with low NUE as below the mean minus one standard deviation (< 0.362). Among these, ten pigs had very high NUE values (mean plus 2 SD; 0.453 to 0.593), all of which had both liver and muscle samples available. Forty-five pigs were classified as having high NUE (0.423-0.450). Of these, three were not sampled, four had only liver samples but lacked muscle tissue, and another four had muscle samples available but missing liver tissue. Ten pigs had very low NUE (mean - 2 SD; 0.265 to 0.331); nine had both tissues available, and one had only a liver sample. Fifty-seven pigs were classified as low NUE (0.337-0.361); among these, no samples were collected from sixteen pigs, two had only liver samples, and three had only muscle tissue.

### RNA isolation

Total RNA was extracted from the muscle samples using the RNAeasy Plus Universal kit (Qiagen, Hilden, Germany). Briefly, 80 mg of muscle tissue was put in 900 *µ*l of QIAzol lysis reagent (Qiagen, Hilden, Germany) in Precellys CK28 tubes (Bertin Technologies SAS, Montigny-le-Bretonneux, France). The mix was then homogenised in a Precellys 24 homogeniser for 2 min at maximum speed. The resulting solution was recovered and processed according to the manufacturer’s instructions. A total of 10 mg of liver was used for total RNA extraction using the RNeasy mini kit (Qiagen, Hilden, Germany). The tissue sample was disrupted and homogenised in 350 *µ*l RLT buffer in Precellys CK14 tubes, and the lysate was recovered after centrifugation and processed according to the manufacturer’s protocol, including the optional DNAse digestion step. For both protocols, RNA was eluted in 40 *µ*l of RNAse-free H_2_O. RNA concentration was determined by fluorometry, using a Qubit quantification system (ThermoFisher Scientific, Waltham MA, USA) and a Quantifluor RNA system (Promega, Madison WI, USA). RNA purity was determined spectrophotometrically (NanoDrop, Thermo Fisher Scientific, USA) by measuring absorbance ratios (A260/A280 and A260/A230) and RNA integrity was assessed with a capillary electrophoresis system, using a QSep 100 (Bioptic, New Taipei City, Taiwan) and high sensitivity RNA cartridge NR1 (Bioptic, New Taipei City, Taiwan). Only samples with an RNA integrity number (RIN) >7 were included in subsequent analyses, as this level of RNA integrity is generally considered suitable for reliable library preparation and sequencing in bulk RNA-seq and was specifically required for the BRB-seq protocol (see below). The RNA samples were aliquoted to avoid repeated freeze–thaw cycles and stored at -80 °C until use for library preparation and sequencing.

### RNA sequencing and creation of the read-count matrix

Samples were split into two balanced pools according to tissue type, NUE level, and sex. Libraries were prepared according to the BRB-seq protocol (Alpern et al., 2019), following the manufacturer’s recommendation (Alithea Genomics). The FASTQ files of the pooled library were generated using the bcl2fastq software (Illumina, v. 2.20.0.422). Mapping and quantification of reads were performed against the Sus scrofa 11.1 reference genome assembly (Warr et al., 2020) using the STAR aligner (v. 2.7.10b, Dobin et al., 2013). The pooled BRB-seq libraries were processed using STARsolo (Kaminow et al., 2021) with the MERCURIUS barcode whitelist, a 14-bp sample barcode and a 14-bp UMI extracted from Read 1, forward-strand specification, and genelevel quantification. Exact command-line parameters (README file) and the barcode whitelist together with the barcode-to-sample mapping are provided in the supplementary information (Monney et al., 2026). The resulting gene count matrices were further processed using custom R scripts to assign barcodes to samples and generate the final count matrix with samples as columns and genes as rows. The generation of libraries, quality assessments, mapping and feature counts were conducted by the Next-Generation Sequencing Platform in collaboration with the Interfaculty Bioinformatics Unit, both from the University of Bern. Sequencing was carried out at Novogene Inc.

### Statistical analysis

Statistical analysis was performed in R (Version 4.5.3, R Core Team, 2025). For quality control, we visually assessed the plots of correlation and principal components analysis of the normalised reads, which showed that the samples of individuals who received a different diet (not reduced in crude protein) differed markedly from the rest of the data. Therefore, we excluded these samples from further analysis. In addition, only genes with at least 3 counts in more than 20 samples were considered for the analysis. Differential gene expression was conducted using the DESeq2 package (version 1.48.1, Love et al., 2014). To define the optimal set of explanatory variables in the model, an iterative model selection (Supplementary Figure S1) was performed on the Agroscope HPC cluster. It was implemented as an iterative tree search starting from the full model including the following explanatory variables: NUE phenotype, sex, slaughter age, weight at birth and at slaughter, ADG, ADFI, FCR, ambient temperature, and age at the start of the experiment. Models were fitted using DESeq2’s maximum likelihood option. At each step, all reduced models obtained by removing one explanatory variable were compared with the full model using gene-wise likelihood ratio tests. Only branches in which fewer than 0.1% of genes showed a significantly better fit for the full model were pursued further; branches failing this criterion were not explored further. To avoid redundant computation, identical reduced models reached through different branches were tested only once. The final model was the simplest model that satisfied the selection criterion. The 0.1% threshold was chosen as a pragmatic cutoff to allow the removal of a variable only when its exclusion significantly worsened the model fit for fewer than 0.1% of genes. For the statistical analysis of differential gene expression, the model with the lowest number of explanatory variables that met the selection criterion was kept to compare groups of individuals with high and low NUE. The DESeq2 independent filtering method was applied after model fitting and prior to DEG testing using a mean normalised count cutoff to exclude lowly expressed genes and enhance detection power without increasing the false discovery rate. A nominal p-value threshold of 0.05 and an adjusted p-value threshold of 0.01 were applied to control for multiple testing. Finally, the log2 fold-change estimates were adjusted using the lfcShrink with the “apeglm” method to reduce noise from low count and/or highly variable genes Zhu et al., 2019. Ensembl gene name were associated to NCBI accession ID and NCBI gene description using biomaRt R package (v. 2.64.0, Smedley et al., 2009) and the Ensembl release 114 (Dyer et al., 2025)

### Pathway analysis

To gain a more comprehensive understanding of our results, we employed a dual approach. We performed over-representation analysis (ORA) to identify pathways significantly enriched among differentially expressed genes (DEGs), and gene-set enrichment analysis (GSEA) to reveal broader, coordinated changes in pathway activity, including subtle shifts that may not reach statistical significance at the single-gene level. Both procedures were performed using the Gene Ontology (GO)(Ashburner et al., 2000; The Gene Ontology Consortium et al., 2023) and the Kyoto Encyclopaedia of Genes and Genomes (KEGG)(Kanehisa et al., 2025; Kanehisa and Goto, 2000) databases as references. GO term analysis provides a hierarchical classification of gene functions while KEGG pathway analysis helps to understand how genes and their products interact within molecular networks. Together these approaches provide a comprehensive understanding of gene function and molecular mechanisms. Genes identified as being differentially expressed were matched with their corresponding Entrez Gene ID and were subsequently used in ORA. In parallel, the genes were ranked according to their p-value and the sign of the log_2_fold-changes obtained from the differential expression analysis to perform GSEA.

The clusterProfiler (v.4.16.0, Wu et al., 2021) and enrichplot (v.1.28.4, Yu, 2025) packages in R were used to hierarchically cluster significantly enriched terms/pathways in a tree. The Jaccard index was used to assess the similarity between the gene sets, followed by the unweighted pair group method with an arithmetic mean (UPGMA, Drummond and Rodrigo, 2000) algorithm for clustering. The number of subgroups to highlight in the tree was determined using a gene-concept network visualisation. This method maps relationships between genes and biological pathways/terms by linking shared genes and spatially separating weakly connected or isolated nodes, which helps identify and prioritise the most biologically coherent and functionally relevant clusters in the dataset.

Data processing and wrangling were conducted using the dplyr package (v.1.2.0, Wickham et al., 2026), and data reshaping was performed using tidyr (v.1.3.2, Wickham et al., 2025), both as part of the tidyverse collection of R packages (v.2.0.0, Wickham et al., 2019). Figures were generated using either the functions available in the aformentioned packages or using ggplot2 (v.4.0.2, Wickham, 2016), with ggrepel employed for improved label placement (v.0.9.7, Slowikowski, 2026), EnhancedVolcano for visualization of differential expression results (v.1.26.0, Blighe et al., 2025), pheatmap for heatmap construction (v.1.0.13, Kolde, 2025), pathview for pathway-based data visualization (v.1.48.0, Luo et al., 2013), and RColorBrewer for color palette implementation (v.1.1-3, Neuwirth, 2022).

## Results

### Sample selection and data quality control

From a larger pool of sampled pigs described in Ewaoluwagbemiga et al. (2023), we selected 48 with the highest and 48 with the lowest NUE, and extracted RNA from their liver and skeletal muscle tissue. After quality assessment of the count matrix Supplementary Figure (S2), three muscle and one liver sample were excluded, leaving 93 muscle and 95 liver samples for RNA sequencing. An additional 14 samples were excluded because the pigs had been fed a different diet (not reduced in crude protein), resulting in 87 samples per tissue type. Of these, 80 pigs contributed samples for both tissues, 7 provided only liver samples, and 7 provided only muscle samples. The final dataset comprised 42 and 41 low-NUE samples and 45 and 46 high-NUE samples for the liver and muscle, respectively (Table 2 and Table 3). Read mapping yielded 35,690 genes in both tissues. Following the removal of features with low counts (<20 in <3 individuals), 13,213 genes in the liver and 12,682 genes in the muscle were retained for differential expression analysis.

**Table 2.**
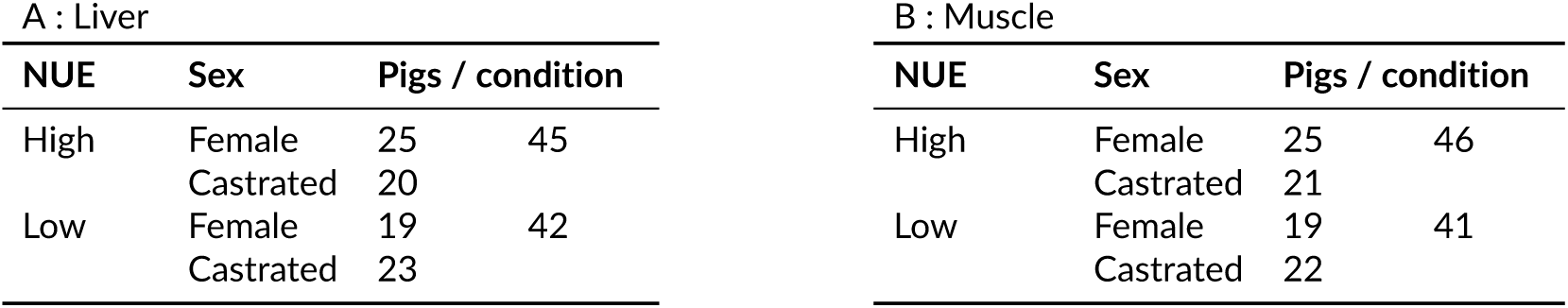
Overview of samples by nitrogen use efficiency (NUE) phenotype, sex, and tissue

| A : Liver |  |  |  | B : Muscle |  |  |  |
| --- | --- | --- | --- | --- | --- | --- | --- |
| NUE | Sex | Pigs / condition |  | NUE | Sex | Pigs / condition |  |
| High | Female | 25 | 45 | High | Female | 25 | 46 |
|  | Castrated | 20 |  |  | Castrated | 21 |  |
| Low | Female | 19 | 42 | Low | Female | 19 | 41 |
|  | Castrated | 23 |  |  | Castrated | 22 |  |

**Table 3.**
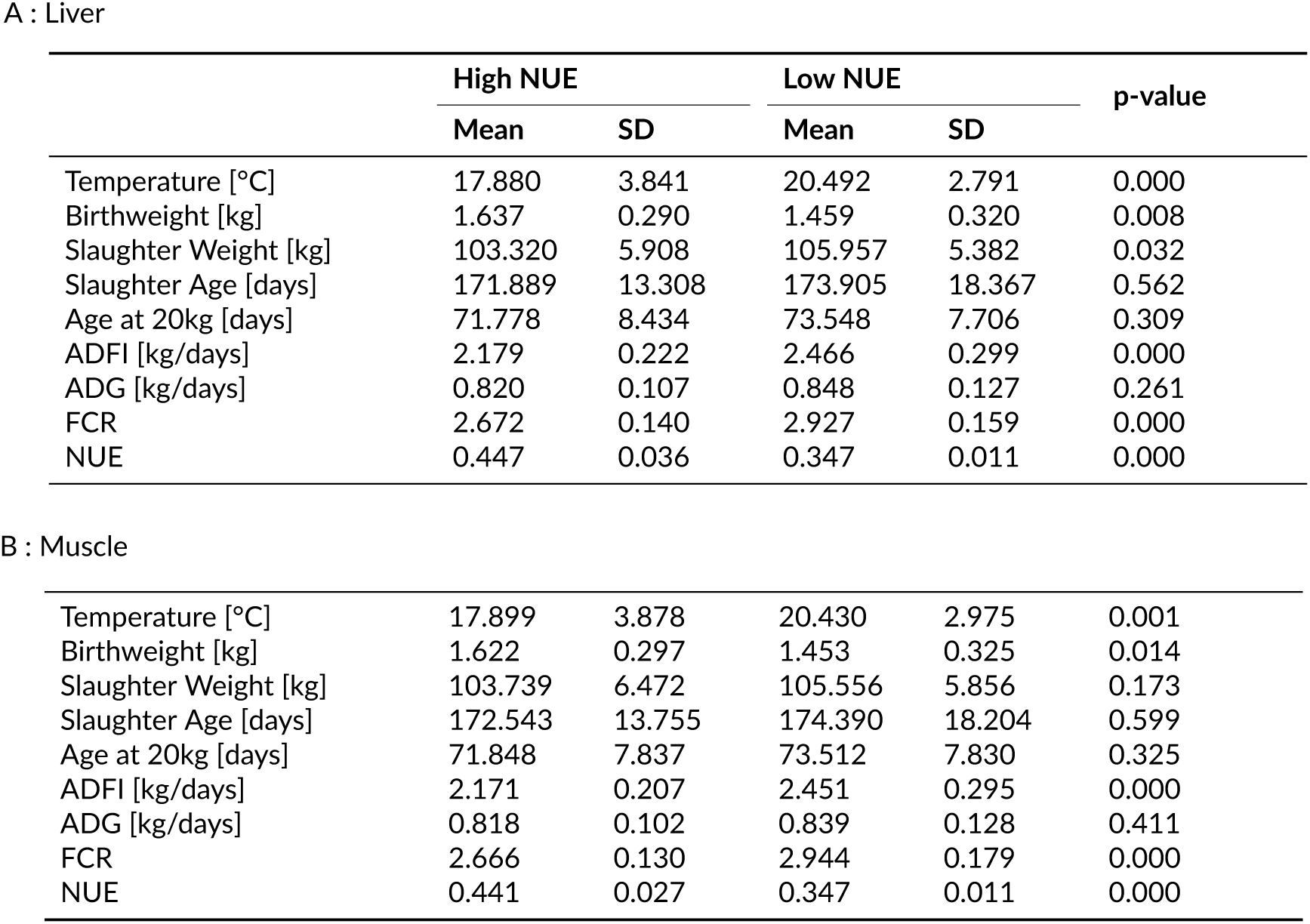
Descriptive statistics and group comparisons for experimental variables in the high and low nitrogen use efficiency groups. Statistical significance between two groups was assessed using a Welch two sample t-test, and the resulting p-values are reported.

A : Liver
|  | High NUE |  | Low NUE |  | p-value |
| --- | --- | --- | --- | --- | --- |
|  | Mean | SD | Mean | SD |  |
| Temperature [°C] | 17.880 | 3.841 | 20.492 | 2.791 | 0.000 |
| Birthweight [kg] | 1.637 | 0.290 | 1.459 | 0.320 | 0.008 |
| Slaughter Weight [kg] | 103.320 | 5.908 | 105.957 | 5.382 | 0.032 |
| Slaughter Age [days] | 171.889 | 13.308 | 173.905 | 18.367 | 0.562 |
| Age at 20kg [days] | 71.778 | 8.434 | 73.548 | 7.706 | 0.309 |
| ADFI [kg/days] | 2.179 | 0.222 | 2.466 | 0.299 | 0.000 |
| ADG [kg/days] | 0.820 | 0.107 | 0.848 | 0.127 | 0.261 |
| FCR | 2.672 | 0.140 | 2.927 | 0.159 | 0.000 |
| NUE | 0.447 | 0.036 | 0.347 | 0.011 | 0.000 |
| B : Muscle |  |  |  |  |  |
| Temperature [°C] | 17.899 | 3.878 | 20.430 | 2.975 | 0.001 |
| Birthweight [kg] | 1.622 | 0.297 | 1.453 | 0.325 | 0.014 |
| Slaughter Weight [kg] | 103.739 | 6.472 | 105.556 | 5.856 | 0.173 |
| Slaughter Age [days] | 172.543 | 13.755 | 174.390 | 18.204 | 0.599 |
| Age at 20kg [days] | 71.848 | 7.837 | 73.512 | 7.830 | 0.325 |
| ADFI [kg/days] | 2.171 | 0.207 | 2.451 | 0.295 | 0.000 |
| ADG [kg/days] | 0.818 | 0.102 | 0.839 | 0.128 | 0.411 |
| FCR | 2.666 | 0.130 | 2.944 | 0.179 | 0.000 |
| NUE | 0.441 | 0.027 | 0.347 | 0.011 | 0.000 |

### Model selection for differential gene expression analysis

Model selection for the differential expression analysis of NUE in the liver identified two models with similar performance to the full model, as determined by a likelihood ratio test (LRT; with 0 rejected genes for LRT against the full model). Both models included the variables ’sex’ and ’temperature’. In addition, one model included ’ADG’ and the other included ’ADFI’. Five equally well-performing models were identified during model selection in the muscle (with 0 rejected genes for LRT against the full model). All five models included ’sex’ and ’ambient barn temperature’ as variables, three included ’age at slaughter’, two included ’ADG’, ’age at 20 kg’ and ’FCR’, and one included ’ADFI’. For consistency of liver and muscle analyses, we selected the models with the most similar parameters for the differential expression analysis:

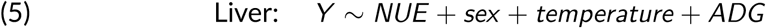

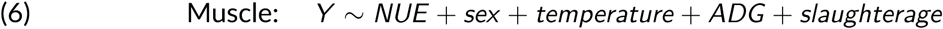

### Differentially expressed genes in pigs with high versus low nitrogen use efficiency

Applying the selected model to each tissue, 177 DEGs were identified in the liver, 54 of which were expressed at higher levels in pigs with high NUE (Fig. 1A and C, Supplementary Table S1). No genes were removed after independent filtering. In skeletal muscle, 133 DEGs were identified, 50 of which were more abundantly expressed in pigs with high NUE compared to those with low NUE (Fig. 1B and D, Supplementary Table S2). In this tissue, 7,622 genes were removed by independent filtering, corresponding to 60.1% of all detected genes. This likely reflects the highly specialised transcriptional profile of this tissue, in which a relatively small set of strongly expressed muscle-related genes dominates overall expression. One gene, *SHMT2*, was differentially expressed in both the liver and muscle. Table 4 lists *SHMT2*, alongside the 10 DEGs with the smallest adjusted p-value and an absolute fold change greater than 2, for each tissue.

**Figure 1.**
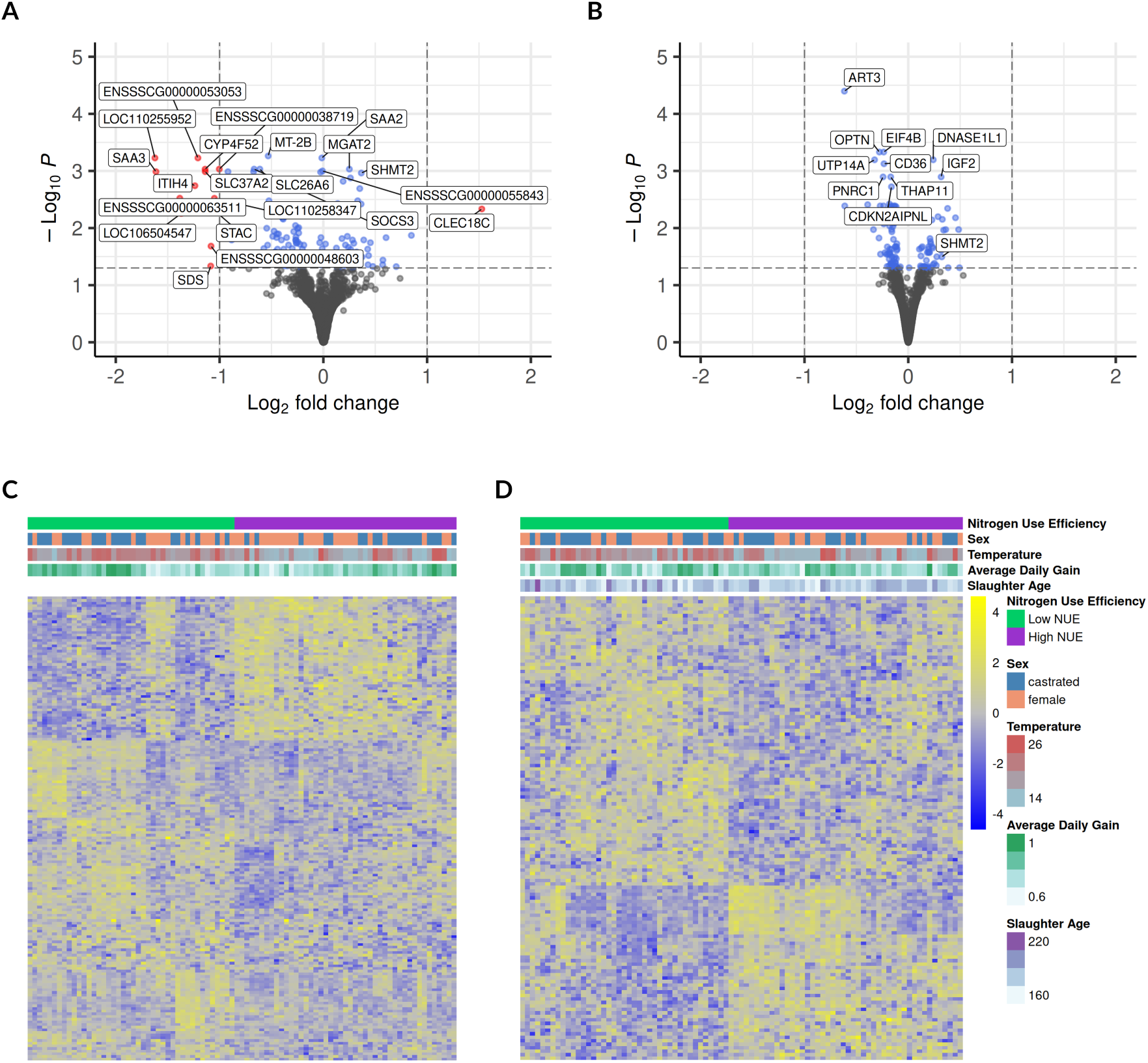
Volcano plots of (A) liver and (B) muscle. The top ten genes with the smallest adjusted p-value and genes with a fold-change larger than 2 are labelled (see Table 4). Heatmap of the differentially expressed genes in liver (C) and skeletal muscle (D). Prior to visualisation, expression values were standardised per gene using Z-score transformation. Colors represent relative expression levels, with positive Z-scores (yellow) indicating higher-than-average expression and negative Z-scores (blue) indicating lower-than-average expression for each gene. The annotation tracks shown above the heatmap represent the covariates included in the statistical model.

**Table 4.**
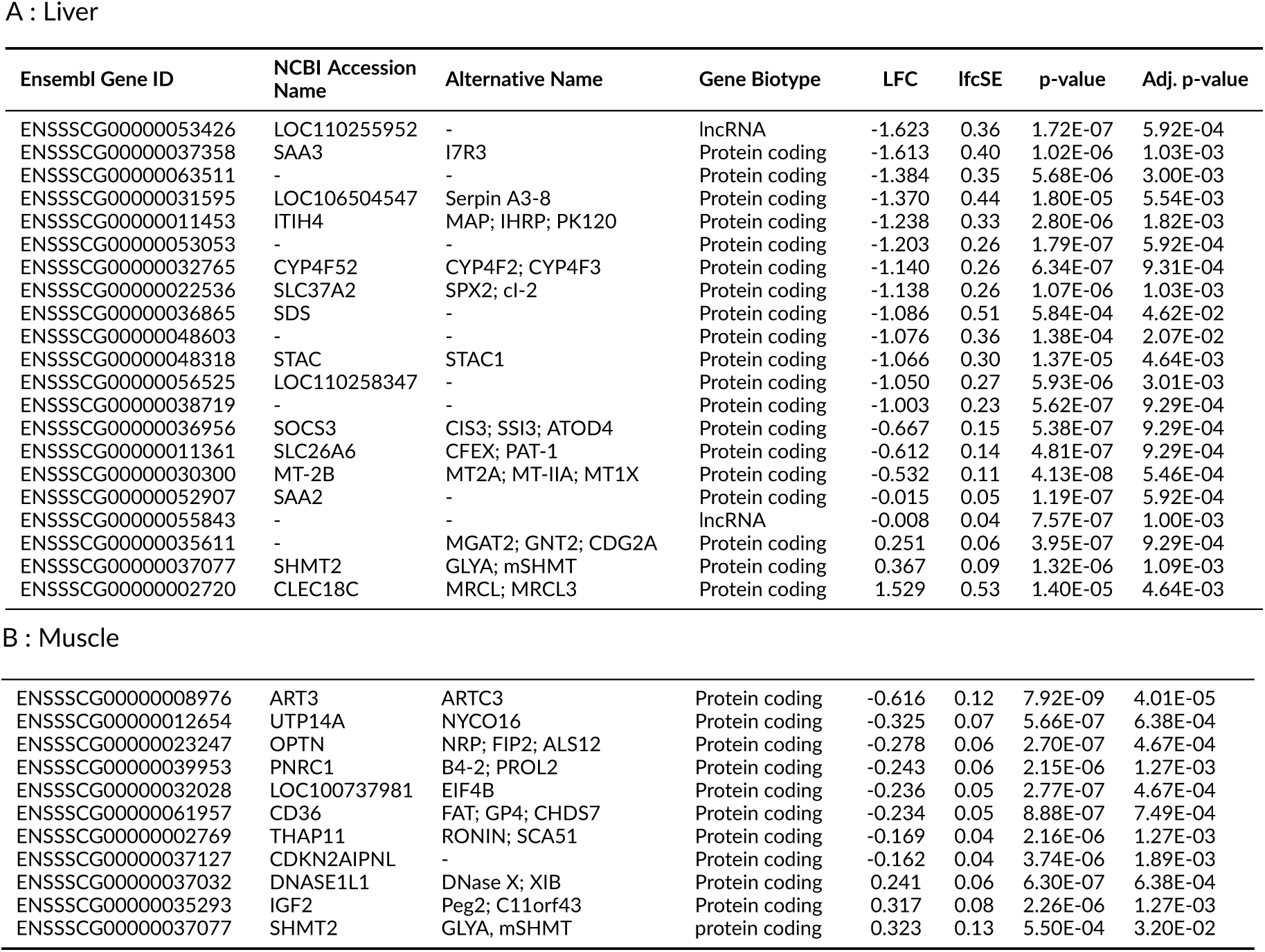
Partial list of differentially expressed genes found in (A) liver and (B) muscle. Only the 10 genes with the smallest adjusted p-value along with the genes with a fold-change larger than 2 are displayed. Genes are ordered according to their fold-change with respect to the low-NUE average expression. A non-exhaustive list of alternative gene names is provided. The full list of differentially expressed genes can be found in Supplementary Table S1 and S2

A : Liver
| Ensembl Gene ID | NCBI Accession Name | Alternative Name | Gene Biotype | LFC | lfcSE | p-value | Adj. p-value |
| --- | --- | --- | --- | --- | --- | --- | --- |
| ENSSSCG00000053426 | LOC110255952 | - | lncRNA | -1.623 | 0.36 | 1.72E-07 | 5.92E-04 |
| ENSSSCG00000037358 | SAA3 | I7R3 | Protein coding | -1.613 | 0.40 | 1.02E-06 | 1.03E-03 |
| ENSSSCG00000063511 | - | - | Protein coding | -1.384 | 0.35 | 5.68E-06 | 3.00E-03 |
| ENSSSCG00000031595 | LOC106504547 | Serpin A3-8 | Protein coding | -1.370 | 0.44 | 1.80E-05 | 5.54E-03 |
| ENSSSCG00000011453 | ITIH4 | MAP; IHRP; PK120 | Protein coding | -1.238 | 0.33 | 2.80E-06 | 1.82E-03 |
| ENSSSCG00000053053 | - | - | Protein coding | -1.203 | 0.26 | 1.79E-07 | 5.92E-04 |
| ENSSSCG00000032765 | CYP4F52 | CYP4F2; CYP4F3 | Protein coding | -1.140 | 0.26 | 6.34E-07 | 9.31E-04 |
| ENSSSCG00000022536 | SLC37A2 | SPX2; cl-2 | Protein coding | -1.138 | 0.26 | 1.07E-06 | 1.03E-03 |
| ENSSSCG00000036865 | SDS | - | Protein coding | -1.086 | 0.51 | 5.84E-04 | 4.62E-02 |
| ENSSSCG00000048603 | - | - | Protein coding | -1.076 | 0.36 | 1.38E-04 | 2.07E-02 |
| ENSSSCG00000048318 | STAC | STAC1 | Protein coding | -1.066 | 0.30 | 1.37E-05 | 4.64E-03 |
| ENSSSCG00000056525 | LOC110258347 | - | Protein coding | -1.050 | 0.27 | 5.93E-06 | 3.01E-03 |
| ENSSSCG00000038719 | - | - | Protein coding | -1.003 | 0.23 | 5.62E-07 | 9.29E-04 |
| ENSSSCG00000036956 | SOCS3 | CIS3; SSI3; ATOD4 | Protein coding | -0.667 | 0.15 | 5.38E-07 | 9.29E-04 |
| ENSSSCG00000011361 | SLC26A6 | CFEX; PAT-1 | Protein coding | -0.612 | 0.14 | 4.81E-07 | 9.29E-04 |
| ENSSSCG00000030300 | MT-2B | MT2A; MT-IIA; MT1X | Protein coding | -0.532 | 0.11 | 4.13E-08 | 5.46E-04 |
| ENSSSCG00000052907 | SAA2 | - | Protein coding | -0.015 | 0.05 | 1.19E-07 | 5.92E-04 |
| ENSSSCG00000055843 | - | - | lncRNA | -0.008 | 0.04 | 7.57E-07 | 1.00E-03 |
| ENSSSCG00000035611 | - | MGAT2; GNT2; CDG2A | Protein coding | 0.251 | 0.06 | 3.95E-07 | 9.29E-04 |
| ENSSSCG00000037077 | SHMT2 | GLYA; mSHMT | Protein coding | 0.367 | 0.09 | 1.32E-06 | 1.09E-03 |
| ENSSSCG00000002720 | CLEC18C | MRCL; MRCL3 | Protein coding | 1.529 | 0.53 | 1.40E-05 | 4.64E-03 |
| B : Muscle |  |  |  |  |  |  |  |
| ENSSSCG00000008976 | ART3 | ARTC3 | Protein coding | -0.616 | 0.12 | 7.92E-09 | 4.01E-05 |
| ENSSSCG00000012654 | UTP14A | NYCO16 | Protein coding | -0.325 | 0.07 | 5.66E-07 | 6.38E-04 |
| ENSSSCG00000023247 | OPTN | NRP; FIP2; ALS12 | Protein coding | -0.278 | 0.06 | 2.70E-07 | 4.67E-04 |
| ENSSSCG00000039953 | PNRC1 | B4-2; PROL2 | Protein coding | -0.243 | 0.06 | 2.15E-06 | 1.27E-03 |
| ENSSSCG00000032028 | LOC100737981 | EIF4B | Protein coding | -0.236 | 0.05 | 2.77E-07 | 4.67E-04 |
| ENSSSCG00000061957 | CD36 | FAT; GP4; CHDS7 | Protein coding | -0.234 | 0.05 | 8.88E-07 | 7.49E-04 |
| ENSSSCG00000002769 | THAP11 | RONIN; SCA51 | Protein coding | -0.169 | 0.04 | 2.16E-06 | 1.27E-03 |
| ENSSSCG00000037127 | CDKN2AIPNL | - | Protein coding | -0.162 | 0.04 | 3.74E-06 | 1.89E-03 |
| ENSSSCG00000037032 | DNASE1L1 | DNase X; XIB | Protein coding | 0.241 | 0.06 | 6.30E-07 | 6.38E-04 |
| ENSSSCG00000035293 | IGF2 | Peg2; C11orf43 | Protein coding | 0.317 | 0.08 | 2.26E-06 | 1.27E-03 |
| ENSSSCG00000037077 | SHMT2 | GLYA, mSHMT | protein coding | 0.323 | 0.13 | 5.50E-04 | 3.20E-02 |

Among the 20 liver genes highlighted in Table 4A, two were identified as lncRNAs (ENSSSCG00000053426 and ENSSSCG00000055843), for which no further functional information was available. Fourteen of the remaining 18 genes have been annotated in NCBI, whereas for the others, putative functional information could be retrieved from UniProt (The UniProt Consortium et al., 2025) and Reactome (Wu and Haw, 2017. Broadly, these genes could be assigned to several non-mutually exclusive functional categories. These included ’amino acid metabolism and energy partitioning’ (*SDS* (Kashii et al., 2005; Ogawa et al., 2006) and *SHMT2* (Ducker and Rabinowitz, 2017)) and ’immune or inflammatory processes’ (*SAA2*, *SAA3*) (Mohanty et al., 2025; Sack, 2018, *ITIH4* (Bhanumathy et al., 2002), ENSSSCG00000031595, annotated as *SERPINA3-8* in UniProt (Baker et al., 2007; Law et al., 2006), *SOCS3* (Carow and Rottenberg, 2014), *CLEC18C* (Tsai et al., 2018), ENSSSCG00000048603, annotated in UniProt as a Sushi domain–containing protein and linked in Reactome to *C4BPA* (Blom et al., 2004), and three homologues of the human immunoglobulin heavy constant gamma gene family or IGHG4-like, namely ENSSSCG00000053053, ENSSSCG00000063511 and ENSSSCG00000056525, while another gene was characterised as an immunoglobulin-like protein (ENSSSCG00000038719). Further categories were ’metabolic homeostasis, detoxification, and cellular protection’ (*MT-2B* (Coyle et al., 2002), *CYP4F52* (Guengerich, 2008; Hardwick, 2008), and *SLC26A6* (Seidler, 2025)); and ’fatty acid processing and energy production’ (*SLC37A2* (Lai et al., 2026) and *MGAT2* (McFie et al., 2022; Shi and Cheng, 2009)). For one gene, *STAC3*, little information was available on its role in the liver. Although the expression of this gene has been documented in the livers of salmon (Bower et al., 2012) and pigs (PigGTEx; Teng et al., 2024; and Bgee; Bastian et al., 2021), its role in hepatic metabolism remains poorly understood. None of the 133 DEGs in skeletal muscle had an absolute fold change greater than 2. All 10 most significant DEGs listed in Table 4B were protein-coding genes. Overall, they could be assigned to several functional categories, ’insulin-related regulation and nutrient deposition’ (*IGF2* (Choi et al., 2025; Jiao et al., 2013), *CD36* (Samovski et al., 2018), *ART3* (Wu et al., 2007), *THAP11* (Hancock et al., 2019)); ’regulation of protein turnover and translation’ (*OPTN* (Kadam et al., 2026), *EIF4B* (Quintas et al., 2024; Walker et al., 2013), *THAP11* (Dejosez et al., 2010), *UTP14A* (Black et al., 2018)); and ’transcriptional or cell-cycle regulation’ (*PNRC1* (Batterson et al., 2023), *THAP11* (Dejosez et al., 2010), *CDKN2AIPNL* (Atrian-Afiani et al., 2023)). Although knockout mice have been reported to show reduced fatigue tolerance during running, the function of *DNASE1L1*, a membrane-anchored enzyme predominantly expressed in skeletal and cardiac cells, remains unclear in this context (Keyel, 2017; Rashedi, 2008).

### Pathway analysis

In total, 14 DEGs in the liver and one in the muscle, including the genes mentioned above, had no matches in the NCBI database; therefore, they were not considered for subsequent pathway analysis. In addition, six liver and two muscle DEGs were excluded from ORA because they corresponded to the same Ensembl IDs. ORA was performed on the remaining 157 liver DEGs, identifying 4 significantly overrepresented KEGG pathways and 27 significantly overrepresented GO terms. The over-representation of KEGG pathways is explained by 13 of the DEGs (Figure 2A and Supplementary Table S3), which were all related to ‘amino acid biosynthesis and metabolism’, particularly serine, cysteine, glycine and glutamate. Of the 27 GO terms, 23 were biological processes and 4 were related to molecular functions; these contained 39 of the DEGs (Figure 2B and Supplementary Table S4). For GSEA, we first sorted the 11,860 annotated genes according to their p-value and the sign of the fold change, considering the low-NUE cohort as the base level. The GSEA highlighted 19 KEGG pathways (8 positively and 11 negatively enriched; Figure 3, Supplementary Figure S3 and Supplementary Table S5) and 61 GO terms (30 up- and 31 down-regulated; Figure 4 and Supplementary Table S6). The four KEGG pathways identified in ORA were also over-represented in the GSEA analysis, alongside three additional pathways that were also related to ’amino acid metabolism’, three related to the ’immune system’, eight to ’human disease’ pathways and one to ’cellular detoxification’, specifically ’lysosome biogenesis’. The human diseases included four pathways for neurodegenerative diseases, while the others were for cardiovascular diseases, infectious diseases, and autoimmune diseases. These pathways cluster in four different groups, driven mainly by changes in oxidative phosphorylation and ribosomal composition for the largest cluster, changes in amino acid metabolism for the second largest and changes in the complement cascade for the third one. The last cluster only include ’lysosome biogenesis’ (Figure 3 and Supplementary Figures S3 and S4). Among the differentially enriched GO terms, 31 were related to biological processes, 25 to cellular components, and 5 to molecular functions. These can be grouped into four main categories: (i) Protein biosynthesis and processing, (ii) amino acid metabolic and catabolic processing, immune and adaptive response, (iii) cellular detoxification, and (iv) cellular anatomical structure arrangement (Figure 4 and Supplementary Figure S5). Nine of the 27 GO terms found in the ORA were not among the differentially represented GO terms in the GSEA. Two of the non-overlapping GO terms (GO:0009072 and GO:0019842) were directly related to ’amino acid processing’ and four (GO:0002526, GO:0006953, GO:0006959 and GO:0060218) were associated with ’immune response’. Two additional GO terms (GO:0006790 and GO:0044272) were linked to both ’amino acid processing’ and ’immune response’. One term (GO:0008514) comprised several solute carriers that are involved in all of the aforementioned processes and functions.

**Figure 2.**
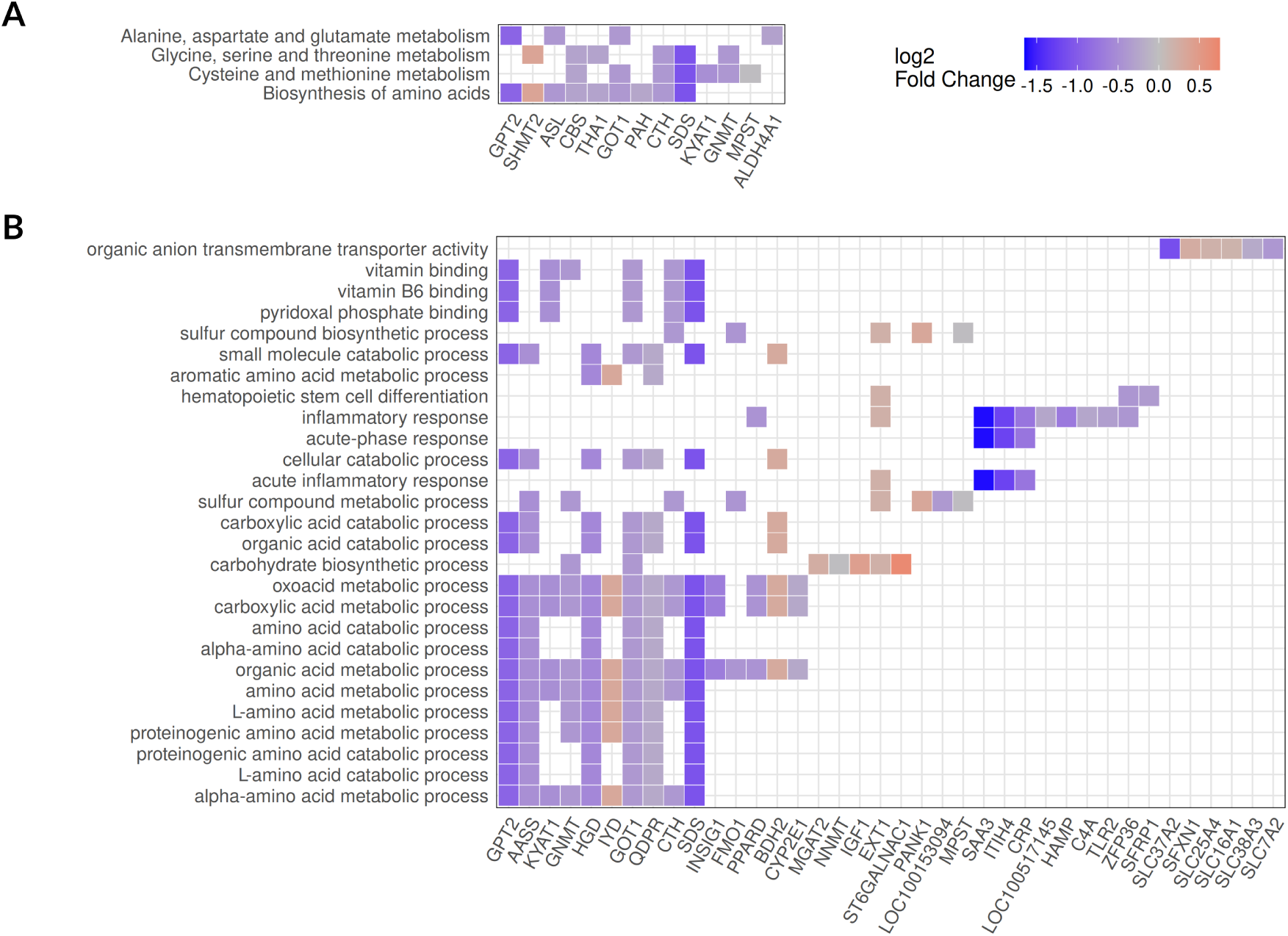
Heatmap of differentially expressed genes in liver identified in over-represented KEGG pathways (A) and GO terms (B). Genes in blue had higher expression in high-NUE pigs compared to low-NUE pigs, whereas genes in red had lower expression. Pathways are are sorted by their adjusted p-value, from bottom to top. Genes are displayed for each pathway, starting from the bottom, first if they are already present in a previous pathway, second according to their adjusted p-value if they are not already on the map. Details on the genes are provided in Supplementary Table S3 and S4.

**Figure 3.**
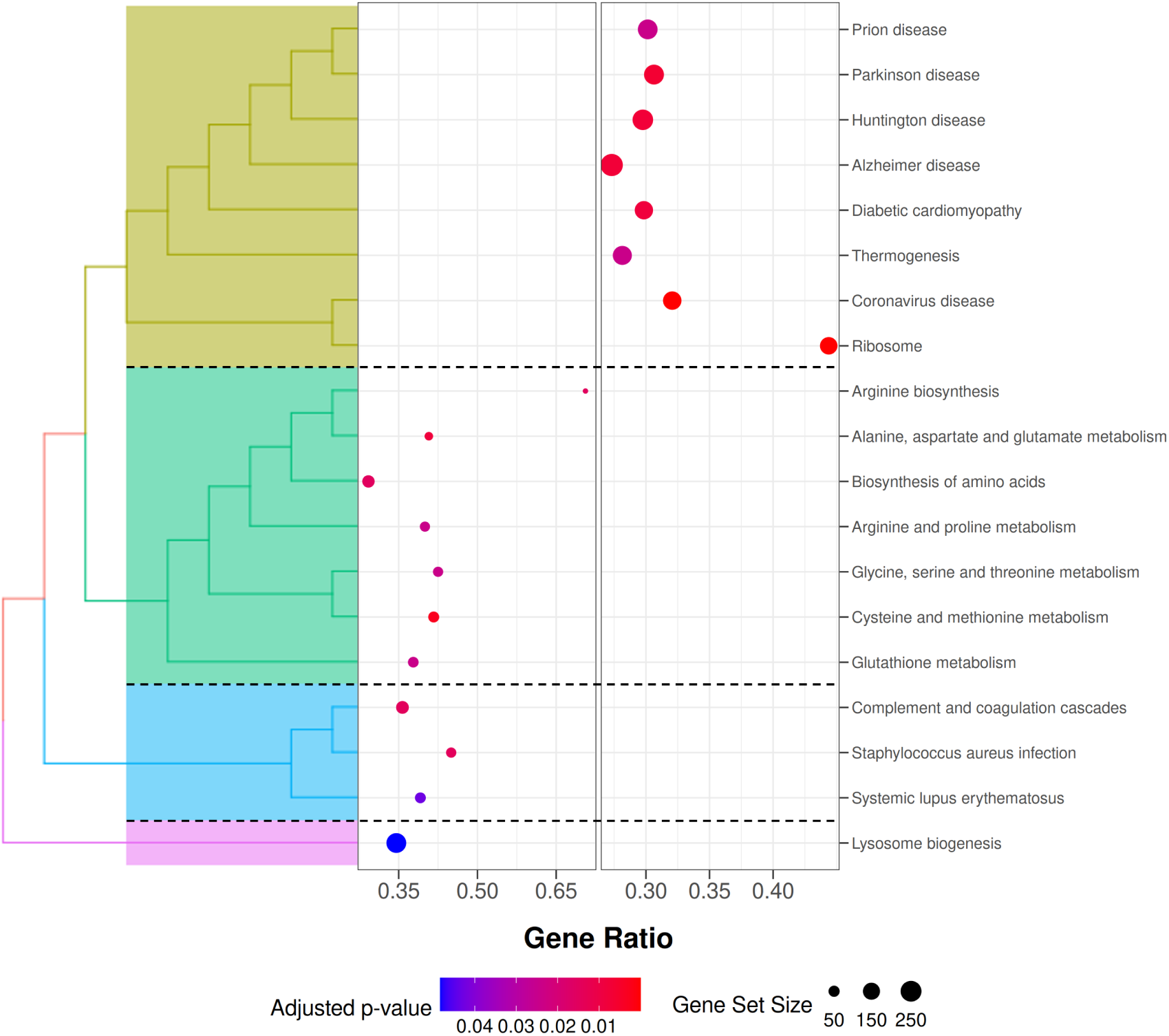
Clustered enriched gene sets identified by GSEA in the liver for KEGG pathways. GeneRatio indicates the proportion of genes in the dataset that contributed to the enrichment, and dot size reflects the absolute number of contributing genes. The colour gradient indicates the adjusted p-value. Genes shown on the left had lower expression overall in high-NUE pigs compared with low-NUE pigs, whereas genes shown on the right had higher expression.

**Figure 4.**
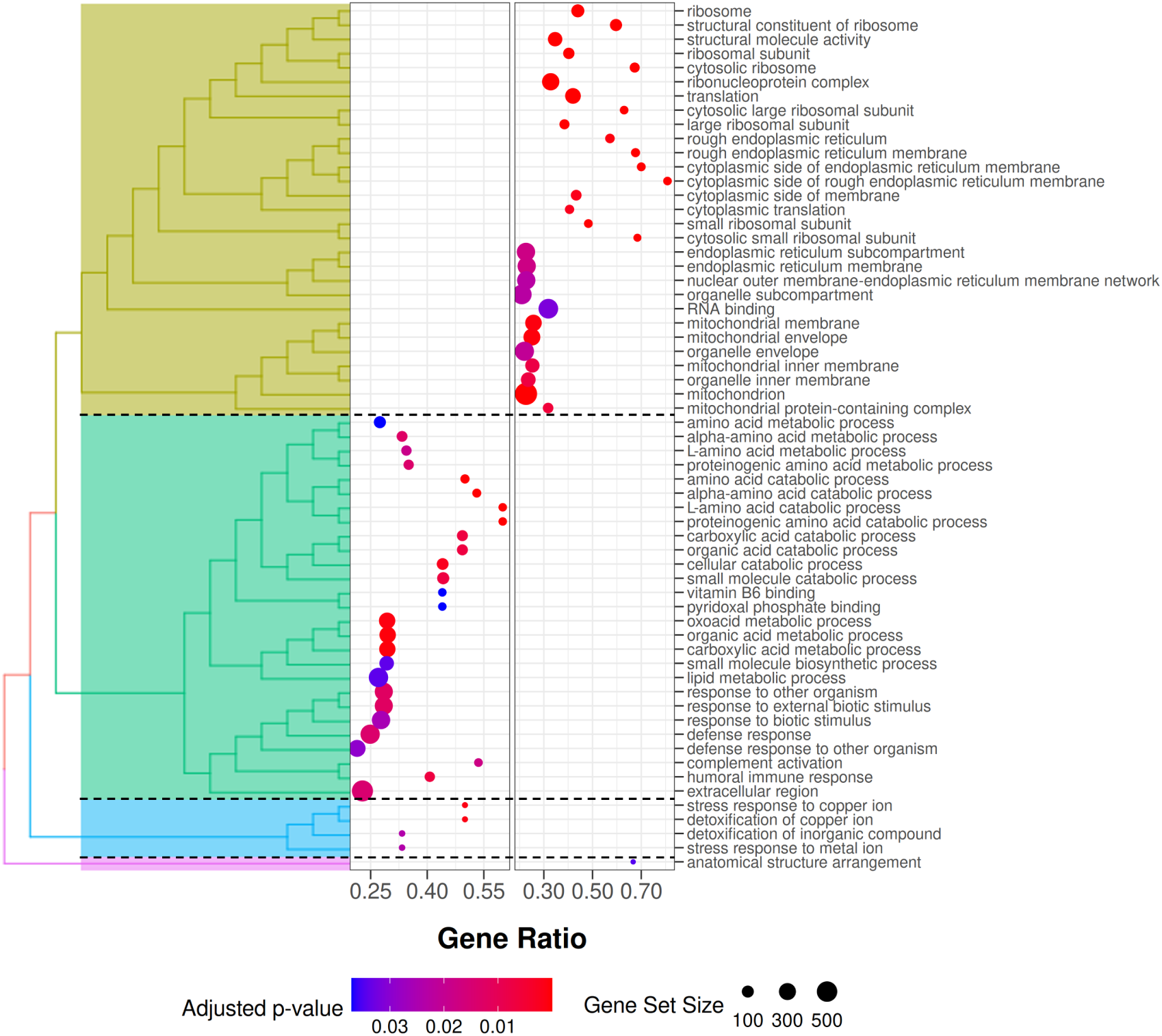
Clustered enriched gene sets identified by GSEA in the liver for GO terms. GeneRatio indicates the proportion of genes in the dataset that contributed to the enrichment, and dot size reflects the absolute number of contributing genes. The colour gradient indicates the adjusted p-value. Genes shown on the left had lower expression overall in high-NUE pigs compared with low-NUE pigs, whereas genes shown on the right had higher expression.

For the muscle samples, ORA was performed on the remaining 130 genes annotated in NCBI. Two KEGG pathways, ‘protein digestion and absorption’ (ssc04974) and ‘aminoacyl-tRNA biosynthesis’ (ssc00970), were found to be significantly over-represented (Supplementary Table S7 and Supplementary Figure S6). However, no significantly overrepresented GO terms were identified.

We performed GSEA on the 4,884 genes identified in the NCBI database, which yielded 13 enriched KEGG pathways, 5 positively and 8 negatively enriched, including the 2 pathways found in the ORA (Figure 5, Supplementary Figure S7 and Supplementary Table S8). These pathways are grouped into two main clusters; one mainly driven by changes in oxidative phosphorylation and one mainly driven by extracellular matrix composition. Two pathways did not belong to these clusters, ‘aminoacyl-tRNA biosynthesis’ (ssc00970) and ’glycine, serine and threonine metabolism’ (ssc00260; Figure 5 and Supplementary Figure S8). The GSEA also yielded 35 GO terms, 9 up- and 26 down-regulated . Sixteen terms were related to biological processes, eighteen to cellular compounds and one to molecular function (Figure 6, and Supplementary Table S9). These could be summarised into five main processes. Three of these encompass several GO terms, ‘respiration and energy production’, ‘matrix structure and morphogenesis process’, and ‘positive regulation of sterol transport’, while the remaining two, ‘serine amino acid biosynthetic process’ and ‘natural killer cell differentiation’, cluster individually (Figure 6, and Supplementary Figure S9).

**Figure 5.**
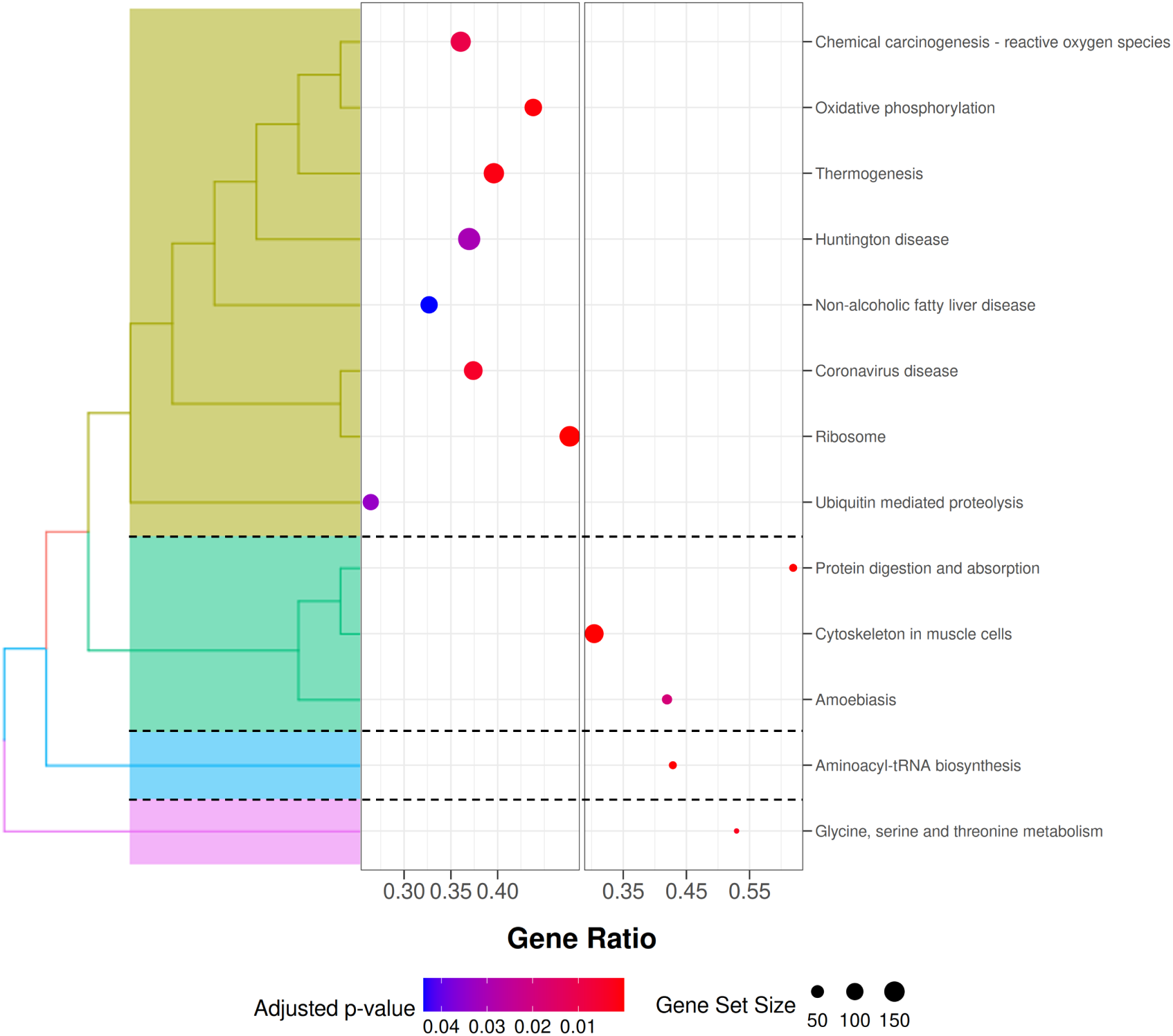
Clustered enriched pathways found in skeletal muscle by GSEA for KEGG pathways. GeneRatio indicates the proportion of genes in the dataset that contributed to the pathway enrichment, while the dot size reflects the absolute number of genes contributing to the enrichment. The colour gradient indicates the adjusted p-value. Left quadrant display gene sets that are overall down-regulated in high-NUE pigs compared to low-NUE pigs while right quadrant display the up-regulated gene sets.

**Figure 6.**
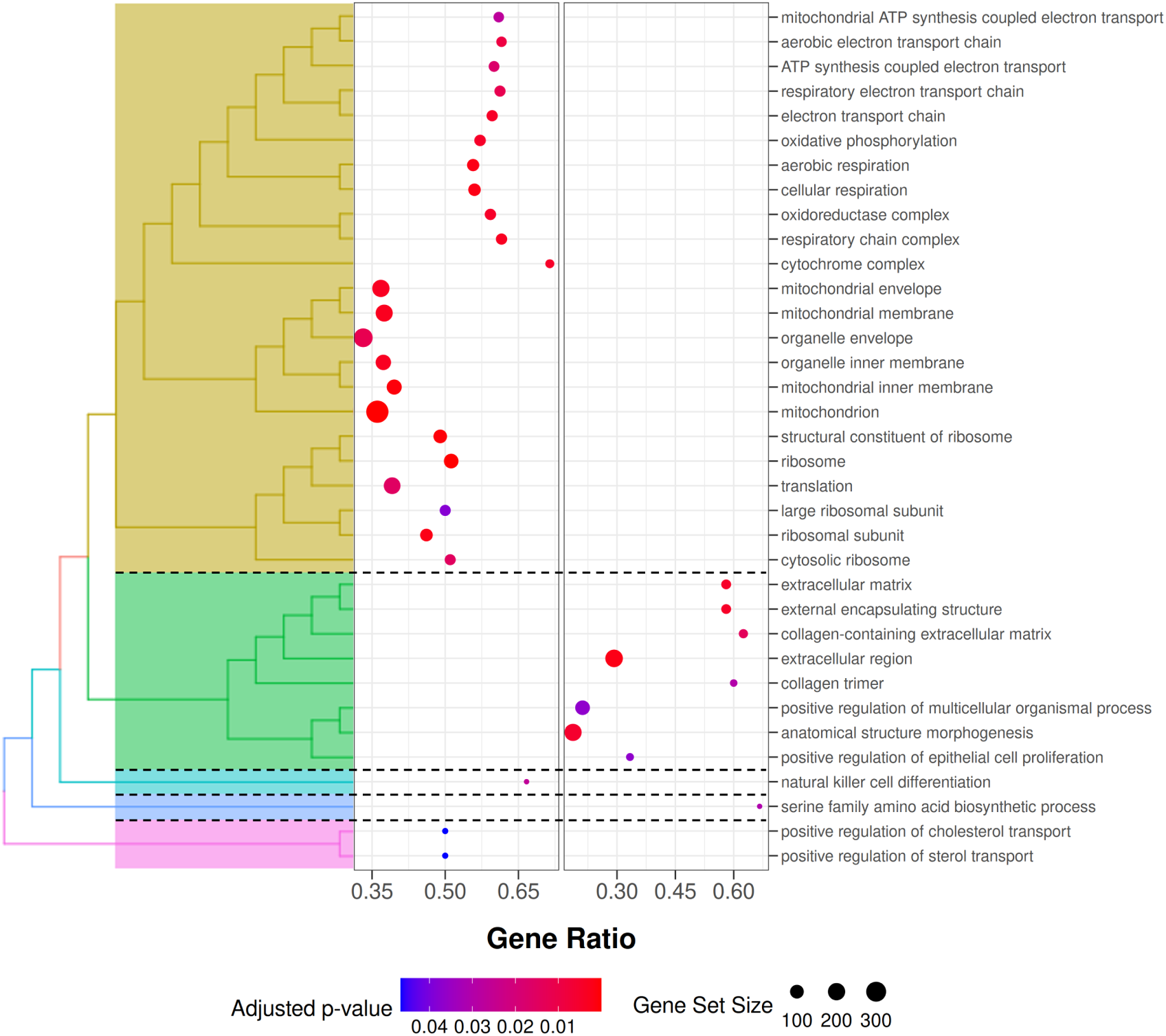
Clustered enriched pathways found in skeletal muscle by GSEA for GO terms. GeneRatio indicates the proportion of genes in the dataset that contributed to the pathway enrichment, while the dot size reflects the absolute number of genes contributing to the enrichment. The colour gradient indicates the adjusted p-value. Left quadrant display gene sets that are overall down-regulated in high-NUE pigs compared to low-NUE pigs while right quadrant display the up-regulated gene sets

## Discussion

In this study, we investigate differences in the gene expression of pigs with high and low NUE in the liver, a central regulator of amino acid metabolism and allocation to key physiological functions, and skeletal muscle, the primary site of protein deposition. The liver and skeletal muscle examined here only capture some of the processes that contribute to variation in NUE. This trait is shaped by a broader system that encompasses nutrient digestion and absorption, endocrine and metabolic regulation, and genetic and environmental factors, among others (Kasper, 2024). Our results show distinct gene expression profiles associated with the NUE phenotype in both liver and muscle tissue, with 177 and 133 DEGs identified, respectively. This suggests that coordinated tissue-specific adaptations play a role in how efficiently nutrients are utilised. To gain insights into the molecular pathways underlying variations in NUE, and to better understand the biological relevance of these gene expression differences, we used pathway enrichment approaches based both on sets of DEGs and on expression patterns across all detected genes in each of the tissues. The results from different pathway detection methods showed substantial agreement and overlap, yielding 18 KEGG pathways and 61 GO terms in the liver and 19 KEGG pathways and 48 GO terms in the muscle. It should be noted that these findings identify transcriptomic differences associated with NUE phenotypes, which may help to shed light on the underlying biological processes of NUE-related variation although they do not, *per se*, establish causal relationships. At the same time, some of these transcriptomic differences may reflect broader efficiency-related physiology, including variation in feed intake, feed conversion, growth performance, or general metabolic state, rather than NUE-specific processes alone. In the following sections, we discuss the findings for each tissue separately before highlighting the shared mechanisms and functional intersections.

### Tissue-specific gene-expression patterns and potential underlying mechanisms

#### Liver

##### Amino acid metabolism and energy partitioning

Both differential gene expression and pathway analyses in the liver suggest key differences between NUE phenotypes in amino acid metabolism and energy partitioning, two processes directly relevant to NUE. In particular, *SDS*, encoding serine dehydratase, showed lower expression in high-NUE pigs. This enzyme catalyses the deamination of serine to pyruvate and ammonia (Kashii et al., 2005; Ogawa et al., 2006). In this reaction, the carbon skeleton of serine is converted into a metabolite that can enter pathways such as gluconeogenesis, while the released nitrogen is subsequently processed through the urea cycle. Regulated by insulin and glucagon (Snell and Walker, 1974), its expression levels correlate negatively with hepatic serine and threonine levels (Joulia et al., 2025). The expression of *SDS* has been shown to respond to dietary protein levels. Higher dietary protein intake increases hepatic SDS expression in rats, likely reflecting the need to metabolise surplus amino acids that exceed the immediate requirements for protein synthesis (Imam et al., 2003). Similarly, increased *SDS* activity has been observed under metabolic conditions associated with accelerated amino acid breakdown, such as fasting or diabetes in rats and mice (Handzlik et al., 2023; SALLACH et al., 1972). Serine levels are likewise reduced in humans with metabolic disorders, including metabolic syndrome and nonalcoholic fatty liver disease (Handzlik et al., 2023; Handzlik and Metallo, 2023; Sim et al., 2020b). Under the protein-restricted conditions of our study, the lower expression of *SDS* in high-NUE pigs may indicate a decreased use of serine as an energy substrate compared to low-NUE pigs.

In both the liver and muscle of the high-NUE pigs, *SHMT2* showed higher expression. It encodes an enzyme that interconverts serine and glycine within mitochondrial one-carbon metabolism (Ducker and Rabinowitz, 2017). In the liver, altered *SHMT2* expression has been linked to lipid metabolism and liver health, although the reported consequences appear context-dependent (Chen et al., 2024; Choi et al., 2022; Wang et al., 2019). Higher *SHMT2* expression in high-NUE pigs may therefore indicate a greater capacity to retain nitrogen in anabolic metabolism rather than excreting it through amino acid catabolism and urea formation. This is further corroborated by the pathway-level results, which show coordinated changes across multiple genes involved in amino acid metabolism and related biosynthetic processes. These changes are consistent with a broader metabolic state that favours the efficient use of nitrogen.

The pathway results confirmed that differences in amino acid metabolism are a central feature distinguishing NUE phenotypes. This signal was largely driven by *SDS* as well as *GOT1*, *GPT2*, and other genes regulating amino acid metabolism from the broader DEG set (Supplementary Tables S3 and S4). In particular, the enriched pathways were related mainly to amino acid metabolism and catabolism, organic acid metabolism, and vitamin B6–dependent enzymatic activity. With the exception of GO:0019842 (vitamin binding), these pathways were also found by GSEA, which also revealed additional amino acid–related processes supported by multiple genes across the ranked gene list (Supplementary Tables S5 and S6). Taken together, these findings indicate that SDS forms part of a broader pattern of coordinated changes across multiple functionally related genes in hepatic amino acid metabolism and homeostasis.

##### Immune function and inflammatory state

Beyond amino acid metabolism, both differential gene expression and pathway analysis also indicated differences in immune function and inflammatory state in the livers of pigs with high and low NUE. Serum amyloid A proteins, such as *SAA2* and *SAA3*, are rapidly and abundantly expressed in the livers of vertebrates during inflammation and infection (Mohanty et al., 2025; Sack, 2018; Uhlar and Whitehead, 1999). While *SAA3* is considered a pseudogene in humans (Sack, 2018), it represents the major acute phase serum amyloid A isoform in pigs (Soler et al., 2011). As SAA proteins are used as biomarkers of inflammation, infection (Eckersall and Bell, 2010) and insulin resistance (Scheja et al., 2008), their lower expression in high-NUE pigs may reflect reduced inflammatory or physiological stress and possibly improved insulin sensitivity and, consequently, more efficient hepatic glucose metabolism. Also expressed at lower levels in high-NUE pigs, *ITIH4* is a liver-specific acute-phase protein that encodes a serine protease inhibitor involved in liver development and regeneration through anti-apoptotic and matrix-stabilising activities (Bhanumathy et al., 2002). *ITIH4* is also used as a biomarker of bacterial infection (Ma et al., 2021) and has been associated with non-alcoholic fatty liver disease (Iguchi et al., 2021; Nakamura et al., 2019). Interestingly, reduced serine protease inhibition decreased activity may also be evident from the lower expression of a predicted *SERPINA3* isoform, *SERPINA3-8* (LOC106504547; *α*-1-antichymotrypsin), in high-NUE pigs. Pigs have several *SERPINA3* family members and isoforms that may have different functions (Irving et al., 2000; Stratil et al., 2002). In general, members of clade A of serpins are involved in inflammatory response where they help limit tissue damage by inhibiting protease activity (Baker et al., 2007; Law et al., 2006). In pigs, hepatic serpin domain-containing proteins have been reported to increase under physiological challenges, including induced sepsis (López-Martínez et al., 2022) and high doses of iron supplementation (Qian et al., 2025). In addition, members of the *SERPINA3* family, including *SERPINA3-8*, differed in serum abundance between pig lines divergently selected for feed efficiency, suggesting potential relevance to efficiency-related physiological states (Grubbs et al., 2016).

Although ORA did not identify any KEGG pathways related to these processes, it highlighted three GO terms, driven mainly by the differential expression of *SAA3*, *ITIH4* and *CRP*, the latter from the broader DEG list (Supplementary Table S1). Additional support came from GSEA, which highlighted the complement and coagulation cascade pathway, involving *ITIH4*, *SERPINA3-8* and *CRP*, as well as several human disease pathways. These enrichments reflected differential expression of genes related to complement activity, inflammatory signalling, and immune regulation, including *NF-κB*, *STAT3*, MHC class II-related immune genes, and the tumor suppressor *p53*. In addition, the signal was influenced by several genes whose effects can be either pro- or anti-inflammatory depending on context, including *PI3K*, *TNFα* and *Wnt*, together with several subunits of oxidative phosphorylation complexes I, III, IV and V. Although some human disease pathways appeared enriched, this does not necessarily imply activation of the corresponding disease pathways. Instead, these signals likely arise from shared immune, inflammatory, and metabolic genes and are therefore more appropriately interpreted in terms of broader functional processes. The interpretation of reduced inflammatory activity is corroborated by the GSEA results for GO terms, which included several terms related to immune function and inflammatory response that showed lower enrichment in high-NUE pigs.

Further support for reduced immune activity and inflammatory signalling comes from the lower expression of the cytokine signalling regulator *SOCS3* in high-NUE pigs. *SOCS3* acts as a negative regulator of cytokine signalling by inhibiting the JAK–STAT pathway (Kershaw et al., 2013), which is activated by cytokines and certain hormones, including Growth Hormone (Liu et al., 2015), and has been suggested to contribute to lower *IGF1* levels through this mechanism (Croker et al., 2008; Kiu and Nicholson, 2012). However, *SOCS3* is also linked to metabolic regulation, especially insulin sensitivity (Torisu et al., 2007), leptin signalling, and energy homeostasis (Liu et al., 2023). In young goats, hepatic *SOCS3* gene expression increases in response to low-protein diets, suggesting that protein restriction may impair somatotropic signalling and limit adaptation to such diets (Firmenich et al., 2020). Lower expression of *SOCS3* in high-NUE pigs may therefore reflect a lower requirement to suppress cytokine signalling, either because of reduced inflammatory activity or, more speculatively, because cytokine signalling is more sensitive. It may also point to differences in insulin responsiveness and hepatic energy regulation. *CLEC18C*, a member of the C-type lectin family, was upregulated in high-NUE pigs. C-type lectins are involved in pathogen sensing and innate immune responses, for example, through the recognition of microbial polysaccharides (Weis et al., 1998). Altered expression of *CLEC* family members has been associated with immune activation in the liver (Tsai et al., 2018). Increased *CLEC18C* expression may therefore reflect differences in innate immune signalling or basal immune activity in high-NUE pigs. Another DEG we found is annotated as a Sushi domain–containing protein and shows homology to members of the human *C4BPA* family. This family is involved in complement regulation and coagulation (Blom et al., 2004); however, the precise function of this gene in pigs is not well understood. In addition to these differences in innate immune signalling, the results also point to differences in adaptive immunity. Among the genes expressed at lower levels in the liver of high-NUE pigs compared to low-NUE pigs were several homologues of the human immunoglobulin heavy constant gamma gene family (Table 4A), which encode IgG class antibodies that are abundant in serum and play a central role in adaptive immunity (Garzón-Ospina and Buitrago, 2022; Vidarsson et al., 2014). The reduced expression of these genes may indicate lower activation of the adaptive immune system and, consequently, reduced allocation of nitrogen towards antibody production in high-NUE pigs.

However, note that even though we excluded clearly unhealthy animals from the study, subclinical differences in health status cannot be ruled out. Such differences, including variations in inflammatory burden or physiological stress, may also have contributed to the observed immune-related expression patterns. In addition, because the present data are based on bulk RNA-seq, the signals detected in liver tissue may not exclusively reflect hepatocyte-intrinsic regulation, but also partly reflect differences in the abundance or transcriptional activity of resident or infiltrating immune cells.

##### Metabolic homeostasis, detoxification and cellular protection

Among the GO terms highlighted by ORA for amino-acid metabolic and catabolic processes and immune response in the liver were sulphur compound metabolic and biosynthetic processes. Sulphur-containing metabolites derived from cysteine and methionine contribute to inflammatory signalling, macrophage activation, glutathione synthesis, and sulfation pathways involved in detoxification and excretion of endogenous and exogenous compounds. In this context, *MT-2B*, a member of the metallothionein family of small cysteine-rich proteins, binds essential and toxic metal ions through its sulphur-rich structure, thereby contributing to cellular metal homeostasis and protection against oxidative stress (Coyle et al., 2002; Klaassen et al., 2009; Palmiter et al., 1992). In pigs, hepatic *MT-2B* gene expression and metallothionein protein abundance increases with dietary zinc levels, indicating a role in zinc homeostasis (Brugger et al., 2014; Martins et al., 2024). In the present study, the lower *MT-2B* expression we observed in high-NUE pigs may therefore reflect differences in hepatic metal homeostasis, or lower oxidative and inflammatory stress.

Pathway analysis also pointed to additional pathways relevant to detoxification. In particular, GSEA identified the KEGG ’lysosome biogenesis’ pathway, driven by lower expression of a broad set of genes encoding, among others, transmembrane proteins involved in lysosomal acidification, proteins mediating Ca2+ export into the cytosol, and several lysosomal acid hydrolases. This pattern suggests reduced lysosomal degradative capacity in high-NUE pigs. GSEA also identified four GO terms related to metal detoxification that were less enriched in high-NUE pigs. Several KEGG pathways related to human diseases similarly suggested shifts in NF*κ*B signalling, oxidative phosphorylation and apoptosis, all closely connected to the production and downstream effects of reactive oxygen species. These are also produced by *CYP4F52*, a pig-specific member of the cytochrome P450 enzyme family that was differentially expressed in our study. Across mammals, *CYP4F* orthologues appear to have broadly conserved functions (Uno et al., 2025). In general, these enzymes oxidise endogenous and exogenous lipid-derived substrates (Guengerich, 2008; Hardwick, 2008), and their expression responds to changes in hepatic substrate load and detoxification demand. Altered expression of *CYP450* family genes has been reported in pigs exposed to cadmium (Wang et al., 2021) and in rats under conditions of elevated nitrogen load (Chung et al., 2002; Guévin et al., 2002; Michaud et al., 2008). The lower expression of *CYP4F52* in high-NUE pigs may therefore reflect a reduced requirement for hepatic oxidative metabolism and detoxification, reflecting lower nitrogen turnover and waste production.

In this context, *SLC26A6* is also of interest. It encodes an anion exchanger involved in ion transport, intracellular pH regulation, and the elimination of metabolic waste products such as oxalate (Seidler, 2025). *SLC26A6* has also been implicated in broader hepatic metabolic responses, including oxidative stress (Remigante et al., 2022) and non-alcoholic fatty liver disease (Chen et al., 2023). In general, solute carrier transporters contribute to metabolic homeostasis and respond to environmental or metabolic challenges, suggesting a role in hepatic detoxification and metabolite transport. Together, these findings fit the broader gene expression pattern observed in this study, which points to a reduced requirement for hepatic stress response and detoxification processes in high-NUE pigs. This includes a lower expression of genes associated with amino acid catabolism (e.g. *SDS*) and inflammatory activity (e.g. *SAA3* and *ITIH4*), suggesting lower production of nitrogenous waste products such as ammonia and urea, as well as a reduced burden of reactive or otherwise potentially harmful metabolic intermediates.

##### Fatty acid processing and energy production

In addition to *SLC26A6*, several other solute carrier genes were differentially expressed in the liver, including genes involved in amino acid transport (*SLC7A2* (Yeramian et al., 2006), *SLC38A3* (Rubio-Aliaga and Wagner, 2016)), energy production (*SLC25A4* (Morrow et al., 2017)), and lactate and pyruvate uptake in hepatocytes (*SLC16A1* (HALESTRAP and PRICE, 1999)). Among these, *SLC37A2* is of particular interest because it showed lower expression in high-NUE pigs and has been linked to glucose metabolism and broader metabolic regulation (Lai et al., 2026). In humans, this gene is associated with glycogen storage disease type Ib, a disorder characterised by impaired glucose homeostasis and excessive glycogen and lipid accumulation in the liver and kidney (Sim et al., 2020a). Variations in *SLC37A2* expression may therefore reflect differences in hepatic energy metabolism between the NUE phenotypes in pigs. This interpretation is supported by previous reports of reduced hepatic activity in glycolytic and other energy-demanding pathways in pigs with high feed efficiency (Xu et al., 2020). Our own pathway results further corroborate this, particularly by GSEA enrichment of GO terms related to mitochondrial function (Supplementary Table S6) and KEGG pathways involving oxidative phosphorylation genes, including subunits of the F-type ATPase complex, with higher expression in high-NUE pigs (Supplementary Table S5).

Another gene consistent with altered hepatic energy partitioning in high-NUE pigs was *MGAT2*, which encodes an enzyme involved in diacylglycerol and triglyceride synthesis, and more broadly in lipid metabolism and energy homeostasis (McFie et al., 2022; Shi and Cheng, 2009). In obese mice, *MGAT2* inactivation improves glucose tolerance and hepatic insulin signalling (Hall et al., 2014). The higher expression of *MGAT2* in high-NUE pigs observed here contradicts reports of reduced hepatic *MGAT2* expression in piglets fed a protein-reduced diet supplemented with monoglyerides and triglycerides (Cui et al., 2022), possibly reflecting differences in developmental stage, dietary context, or metabolic adaptation. In the present study, higher *MGAT2* expression may indicate enhanced lipid processing and energy partitioning, potentially reducing reliance on amino acids as an energy source. This interpretation is supported by the lower *SDS* expression and broader pathway-level signals, suggesting a shift in hepatic energy metabolism away from amino acid oxidation in high-NUE pigs.

##### Cellular structural organisation

In addition to the processes discussed above, the GSEA of both GO terms and KEGG pathways revealed differences in cellular structural organisation. The KEGG results showed lower expression of several tubulins and higher expression of several ribosomal subunits across multiple human disease pathways in high-NUE pigs. This pattern was even more prominent in the GO term analysis, accounting for almost half of the enriched terms, and included terms not only associated with mitochondria, which may be partly related to the shift in energy metabolism, but also with ribosomes, RNA binding and endoplasmic reticulum compartments. Together, these findings suggest broader differences in cellular organisation and translation-related processes. However, as these changes are involved in complex, interconnected mechanisms, it is not possible to confidently infer their functional consequences from the present data alone.

##### Regulation of insulin

Taken together, several of the processes discussed above—namely inflammatory and complement-related signalling, amino acid catabolism, and lipid and energy metabolism—converge on insulin-related regulation as a potential integrative axis that differentiates NUE phenotypes. This connection is supported not only by some of the genes discussed in detail above, which were among the most strongly differentially expressed (Table 4A), but also by additional DEGs (Supplementary Table S1) that did not meet the criteria for inclusion in Table 4A but fit the same broader pattern. Although insulin signalling itself was not directly highlighted in the pathway analysis, several gene- and pathway-level results suggest that it contributes to the observed patterns. Furthermore, many of the DEGs represented in the enriched ORA categories are known to be modulated by insulin or to influence insulin-related processes. This is particularly evident for amino acid catabolism, as insulin suppresses amino acid breakdown and promotes protein synthesis, which is consistent with the lower expression of *SDS* in high-NUE pigs, as discussed in detail above. Additional genes in the broader DEG list supporting this interpretation include *AASS*, *GNMT*, and *KYAT1* (Supplementary Table S1, (Liao et al., 2016; Muñoz et al., 2026; Zhen et al., 2022), all of which participate in different aspects of amino acid metabolism. This link may also extend to inflammatory regulation through *KYAT1*, as altered kynurenic acid production can affect anti-inflammatory signalling (Zhen et al., 2022), in line with the lower expression of *SAA2*, *SAA3*, and *SOCS3*. Although insulin does not directly affect the complement cascade, the variation in inflammatory activity between the NUE phenotypes discussed above, including those relating to the complement cascade, may still partly reflect insulin-related regulation. One example is *C3*, encoded by LOC100517145 in the broader DEG set, which is often present at higher levels under insulin-resistant conditions (Muscari et al., 2007). Its lower expression in high-NUE pigs is, therefore, consistent with more effective insulin action. Finally, insulin also promotes the conversion of excess carbohydrates and fatty acids into triglycerides for storage, a process in which *MGAT2* plays an important role (Cheng et al., 2022). In this context, the higher expression of *MGAT2*, as detailed above, as well as *IGF1* and *INSIG1* (Figure 2A and B and Supplementary Tables S1 and S3), supports the interpretation that high-NUE pigs show a metabolic state that favours lipid processing and storage, and reduces reliance on amino acids for energy production.

#### Muscle

##### Insulin-related regulation and nutrient deposition

As in the liver, insulin-related regulation appears to be a central feature distinguishing the muscle transcriptomes of high- and low-NUE pigs, as several of the genes with the most pronounced differential expression in Table 4B are linked to insulin signalling or related processes. One example is the higher expression of insulin-like growth factor 2 (*IGF2*) in high-NUE pigs. *IGF2* is a hormone structurally related to insulin that can activate the same signalling pathways through binding to its own receptor, *IGF2R*, or to the insulin receptor (Choi et al., 2025; Jiao et al., 2013). It is involved in glucose metabolism in adipose tissue, skeletal muscle, and liver, and plays an important role in muscle cell proliferation and differentiation in mammals (Gerrard et al., 1998). The transcription of *IGF2* is controlled by multiple promoters, some of which are imprinted so that only the paternally inherited allele is expressed, while the maternally inherited allele is silenced (Li* et al., 2008). In pigs, a single-base substitution within a regulatory element of intron 3 (IGF2-G3072A) of the paternally inherited allele increases *IGF2* expression (Aslan et al., 2012) and has been shown to enhance muscle growth and lean meat percentage and to reduce overall fat deposition in pigs (Burgos et al., 2012; Clark et al., 2014; Van Laere et al., 2003).

Further support for differences in lipid uptake and intramuscular fat deposition comes from the lower expression of *CD36* and *ART3* in high-NUE pigs. *CD36* encodes a fatty acid translocase that mediates the uptake of long-chain fatty acids into skeletal muscle cells (Glatz & Luiken, 2017) and has been associated with muscle triglyceride content (Bonen et al., 2004). Beyond its role in fatty acid uptake, *CD36* has also been linked to insulin signalling through its effects on the tyrosine phosphorylation of the insulin receptor (Samovski et al., 2018). In pigs, *CD36* expression was higher in the small intestine and muscle in piglets with low birth weight that were fed a high-fat diet, a pattern thought to contribute to the greater accumulation of triglycerides observed in their muscle tissue (Wang et al., 2023). *ART3*, an insulin-responsive gene encoding ADP-ribosyltransferase (Wu et al., 2007), has also been linked to fat deposition in skeletal muscles during development. For example, in Wannanhua pigs, the intramuscular fat content in the longissimus dorsi muscle increased substantially between 120 and 240 days of age, while the expression of *ART3* in this muscle decreased over the same period (Li et al., 2024). Moreover, in a breed comparison, chromatin accessibility, transcriptional levels, and protein abundance of *ART3* were higher in Leixiang pigs with high intramuscular fat content than in Songliao black pigs with lower intramuscular fat content (Zhang et al., 2025). Thus, the lower expression of these genes in high-NUE pigs in the current study supports the interpretation of reduced fatty acid uptake into skeletal muscle and lower propensity for intramuscular lipid accumulation. However, the intramuscular fat content did not differ significantly between high- and low-NUE pigs in the subset with available data (for 53 animals, approximately 60% of all pigs with muscle samples), averaging 46.58 *±* 13.49 g/kg and 52.22 *±* 21.11 g/kg, respectively (p = 0.242, Welch two-sample t-test; results not shown). This may reflect the limited sample size of the subset or the fact that transcriptomic differences in lipid metabolism and transport do not necessarily translate directly into detectable differences in intramuscular fat at this level. Nevertheless, in the same pig cohort analysed at larger scale, NUE and intramuscular fat showed a moderate negative genetic correlation (-0.39*±* 0.15; Ewaoluwagbemiga et al., 2023), supporting the view that improved NUE may generally be associated with reduced intramuscular fat deposition at the genetic level.

Another gene linked to insulin-related regulation and showing higher expression in high-NUE pigs was *THAP11*. It encodes a transcription factor known to form a complex with HCF-1 and the insulin receptor in the nucleus, thereby enhancing promoter activity (Hancock et al., 2019). Although its specific role in skeletal muscle remains poorly characterised, *THAP11* is known to regulate genes involved in protein biosynthesis and energy production, including several ribosomal subunits and *SHMT2* (Dejosez et al., 2010). *SHMT2* also showed higher expression in the muscles of high-NUE pigs, consistent with the pattern observed in the liver. In hypertrophy- stimulated murine muscle cells, *SHMT2* gene expression increases in response to IGF-related signalling (Baumert et al., 2024). The higher expression observed in our study may indicate an increase in the activity of mitochondrial serine and glycine metabolism in muscle tissue, which could support amino-acid homeostasis and anabolic processes associated with muscle growth.

Pathway analyses provided indirect support for the interpretation that skeletal muscle in high-NUE pigs relies less on amino acid oxidation and more on the broader mitochondrial energy metabolism of non-nitrogen substrates to meet energetic demands. Although ORA did not identify pathways specifically related to lipid uptake or intramuscular fat deposition, GSEA highlighted processes related to mitochondrial energy metabolism, including oxidative phosphorylation and thermogenesis, as well as glycine, serine and threonine metabolism. The enrichment of these pathways in high-NUE pigs suggests that their skeletal muscle metabolism relies more strongly on carbohydrates and lipids for energy production, thereby reducing their reliance on amino acid oxidation.

##### Regulation of protein turnover and translation

Beyond insulin-related regulation, downstream mTOR signalling also appears to contribute to the differences in NUE, given its central role in regulating protein synthesis (Bodine, 2022). Initial support for this comes from the enrichment of the ’protein digestion and absorption’ pathway in ORA in high-NUE pigs. However, the enrichment of this pathway appears to be driven by the differential expression of several collagen subunits, potentially reflecting differences in muscle growth or structure rather than protein digestion *per se*. This interpretation is further supported by the GSEA results, which pointed to differences in both intracellular and extracellular organisation. Together with the change in lipid deposition and in protein synthesis and turnover, these findings suggest that skeletal muscle high-NUE pigs might be more strongly oriented towards growth, whereas low-NUE pigs may be more oriented towards homeostasis. In addition to collagen subunits, this pathway likely also reflected the higher expression of SLC3A2 in high-NUE pigs (Supplementary Figure S7 and Supplementary Table S8). *SLC3A2* encodes the heavy chain of a heteromeric amino acid transporter that associates with *LAT1* to mediate leucine uptake in exchange for glutamine. Given that intracellular leucine accumulation triggers mTOR complex 1 (*mTORC1*) activation, this pattern is consistent with increased signalling towards protein synthesis and muscle growth in high-NUE pigs (Kahlhofer and Teis, 2023).

Several other genes that were differentially expressed between the two NUE phenotypes were related to *mTOR* signalling and protein turnover in skeletal muscle. Notably, *OPTN* showed lower expression in high-NUE pigs. *OPTN* has been identified as a key regulator of the AKT/mTOR pathway through its interaction with RICTOR, the defining component of mTOR complex 2 (*mTORC2*), which influences spatial organisation and cell survival (Kadam et al., 2026). In mice, *OPTN* knockdown delays muscle regeneration and impairs myoblast differentiation, whereas its overexpression promotes myoblast differentiation (Shi et al., 2022). In neonatal pigs, *OPTN* expression has been positively associated with a gene network related to protein degradation (Manjarín et al., 2020). A recent mouse study identified it as a regulator of muscle homeostasis during muscle atrophy, a condition characterised by excessive protein breakdown and reduced protein synthesis (Shi et al., 2026). More generally, *OPTN* participates in selective autophagy and muscle remodelling processes involved in protein turnover (Ying and Yue, 2016). Its lower expression in high-NUE pigs may therefore indicate lower activity of pathways associated with protein turnover. *EIF4B* also showed lower expression in high-NUE pigs. As part of the eukaryotic translation initiation machinery, it promotes protein synthesis by facilitating the recruitment of mRNA to ribosomes (Quintas et al., 2024), as well as by stimulating a helicase enzyme that unwinds folded regions of mRNA, thereby enabling ribosomes to access it and initiate translation (Walker et al., 2013). Its phosphorylation and activity are regulated through convergent signalling from the mTOR and MAPK pathways (Shahbazian et al., 2006). In Wujin pigs, *EIF4B* expression increased in response to high-protein diets at later growth stages and was associated with a significantly higher carcass muscle weight at the same body weight (Liu et al., 2014). More generally, a reduced abundance of components of the eIF4 translation initiation machinery has been associated with reduced muscle hypertrophy in low-birth-weight piglets (El-Kadi et al., 2018). The lower expression observed here may therefore reflect differences in translational regulation between the NUE phenotypes. Together with the *OPTN* results, this suggests that high- and low-NUE pigs differ in the regulation of protein turnover and translation in skeletal muscle, which potentially reflects variations in the balance between protein synthesis and degradation.

Balanced modification of protein synthesis may also involve changes in ribosome biogenesis. As noted above, *THAP11* regulates the expression of several ribosomal subunits, and *UTP14B*, which showed lower expression in high-NUE pigs, is required during an early stage of ribosome assembly (Black et al., 2018). This interpretation is further supported by the enrichment of the KEGG ribosome pathway and six GO terms related to translation and ribosome composition and localisation. Similarly, the enrichment of the aminoacyl-tRNA biosynthesis pathway in ORA further supports differences in translation-related processes between NUE phenotypes. Its enrichment was driven by the higher expression of *AARS1*, *GARS1*, *IARS1*, *MARS1*, and *TARS1* in high-NUE pigs compared to low-NUE pigs (Supplementary Figure S6 and Supplementary Table S7). All five genes encode cytosolic aminoacyl-tRNA synthetases, which attach amino acids to their cognate tRNA and thereby provide essential substrates for protein translation (Gomez and Ibba, 2020). The results of GSEA suggested that this pattern was not restricted to these five genes but extended more broadly across this functional group. GSEA further highlighted pathways related to translation, such as the ribosome pathway and the structural constituent of the ribosome pathway, as well as pathways related to protein turnover, including ubiquitin-mediated proteolysis. These enrichments were driven predominantly by the overall lower expression of genes encoding ribosomal proteins and components of the ubiquitin–proteasome system in high-NUE pigs, rather than any of the DEGs discussed above, suggesting broader coordinated differences in the machinery of protein synthesis and degradation between NUE phenotypes.

##### Transcriptional and cell cycle regulation

Finally, several genes showing lower expression in high-NUE pigs were related to transcriptional regulation and cell cycle–related processes, including *PNRC1*, *CDKN2AIPNL* and the aforementioned *THAP11*. *PNRC1* has been reported to respond to physiological changes in muscle tissue and was reduced in the skeletal muscle of previously sedentary but otherwise healthy lean humans following high-intensity interval training (Batterson et al., 2023). In cattle, *PNRC1* was also identified as a hub gene in a co-expression network associated with intramuscular fat content, where it showed lower expression in entire males than in castrated animals with higher intramuscular fat levels (Reis et al., 2024). *CDKN2AIPNL* has been identified as a candidate gene in genomic regions under selection in pigs (Atrian-Afiani et al., 2023), as a candidate gene for intramuscular fat in Canadian crossbred pigs (Mozduri et al., 2025). A cis-eQTL affecting its expression in skeletal muscle has also been reported in pigs, where this gene has been associated with variations in loin eye size (Steibel et al., 2014). Together, these findings point to broader differences in transcriptional and cell cycle–related regulation between NUE phenotypes, although their specific relevance to NUE remains unclear.

#### Tissue-specific effects on independent filtering

The large number of genes removed by independent filtering in skeletal muscle compared to the liver is likely a consequence of the underlying biology of the tissue rather than a limitation of the analysis. Skeletal muscle is a highly specialised, terminally differentiated tissue in which transcription is dominated by genes related to contraction and other core muscle functions, while many other genes are expressed at low levels (Lindskog et al., 2015). This leads to a comparatively high proportion of genes with low mean counts, which contributes little power to differential expression analysis. Independent filtering in DESeq2 is designed to remove such genes from multiple testing, thereby improving the detection of biologically meaningful differential expression while maintaining an appropriate false discovery rate control (Bourgon et al., 2010; Love et al., 2014). In this context, the filtering of around 60% of the genes in the muscle dataset is therefore not unexpected.

#### Future directions

Future work should examine the associations observed here in independent pig cohorts to assess their robustness and generalisability. In particular, candidate genes such as *SDS*, *SHMT2*, *SAA3*, *SOCS3*, *IGF2*, and *CD36* could be examined in new cohorts under protein-restricted conditions, either through targeted assays such as qPCR or within additional RNA-seq datasets generated from related experiments addressing complementary questions. Because NUE is a complex trait shaped by a broader biological system involving processes beyond liver and skeletal muscle (Kasper, 2024), such studies could also help place the present transcriptomic findings into a wider physiological context by extending analyses to other relevant tissues and levels of biological organisation. Such studies could also test whether these associations are stable across developmental stages and body weights, and extend analyses to additional tissues relevant to NUE, such as intestine, kidney, immune tissues and brain. Where available through public repositories or consortium datasets (Teng et al., 2024), integration with epigenomic or chromatin-level data could provide additional insights into the regulatory architecture underlying these transcriptional differences. More broadly, a multi-omics framework incorporating transcriptomic, proteomic, metabolomic, and genomic variation would help distinguish regulatory mechanisms from downstream physiological consequences (Fang et al., 2025; Kreitmaier et al., 2023; Vandereyken et al., 2023Kreitmaier et al., 2023; Vandereyken et al., 2023; Fang et al., 2025). This could include assessing overlap between differentially expressed genes, previously implicated genomic regions, and convergent biological pathways, as well as integrating circulating biomarkers such as plasma metabolites, urea, amino acids, and acute-phase proteins to link tissue-level transcriptional patterns to systemic physiology. Finally, comparative analyses with other mammals may help establish which of the identified mechanisms are specific to pigs and which reflect more general biological responses to limited protein supply. Together, such approaches would help connect the present RNA-seq findings more directly to the genetic architecture of NUE and clarify which molecular signals are most robust, generalisable, and potentially useful as biomarkers or mechanistic indicators of NUE.

## Conclusion

Overall, differential gene expression and pathway analyses of the liver indicate that high-NUE pigs are characterised by a coordinated metabolic state involving reduced amino acid catabolism. Gene- and pathway-level patterns support this by suggesting that the livers of high-NUE pigs rely more on carbohydrates and lipids than on amino acids for energy, thereby reducing the formation of nitrogen waste and preserving amino acids for productive use. Consequently, non-nitrogen substrates may be more efficiently directed towards energy production and storage pathways, while amino acids are spared from oxidation and retained for protein deposition. Reduced amino acid catabolism may also contribute to lower production of nitrogenous waste products and other potentially harmful metabolic intermediates, consistent with the broader hepatic expression pattern of lower levels of genes involved in detoxification and inflammation. The lower expression of genes related to inflammatory and detoxification processes in the livers of high-NUE pigs may therefore be better interpreted as a consequence of a metabolic state characterised by reduced amino acid catabolism than as evidence that these processes directly drive improved NUE. In such a metabolic state, the decreased production of nitrogenous waste products and other potentially harmful intermediates would be expected to lower the need for hepatic stress response and detoxification activity. In skeletal muscle, the transcriptional profile of high-NUE pigs in indicates reduced lipid uptake, lower reliance on amino acid oxidation, and a greater emphasis on growth-related processes, translational regulation, and protein accretion. These patterns suggest that amino acids are used more efficiently for muscle protein synthesis than for energy production, while energetic demands are increasingly being met through carbohydrates and lipids. Concurrently, the reduced expression of genes associated with fatty acid uptake suggests that lipids are less readily incorporated into muscle tissue, thereby limiting the deposition of intramuscular fat and favouring the accretion of lean tissue. Taken together, these observations suggest that variations in NUE are associated with coordinated, tissue-specific differences in nutrient partitioning, with the liver and skeletal muscle each contributing distinct but complementary aspects of a broader metabolic organisation. In high-NUE pigs, this organisation is consistent with more efficient allocation of nitrogen towards lean tissue growth while energetic demands are increasingly met through non-nitrogen substrates. These transcriptomic patterns provide insight into biological processes associated with NUE variation under protein-restricted conditions, although they do not by themselves distinguish causal drivers from down- stream physiological consequences. The present results help to define the liver- and muscle-associated components of NUE-related biology. Furthermore, they emphasise that variation in NUE originates from the broader integration of the whole animal rather than from these tissues alone.

## Appendix A. Supplementary Tables

Supplementary tables are available online (https://doi.org/10.5281/zenodo.20527211).

Supplementary Table S1: List of genes expressed in the liver, ordered by adjusted p-value. The first 177 genes are significantly differentially expressed (p < 0.05).

Supplementary Table S2: List of genes expressed in skeletal muscle, ordered by adjusted p-value. The first 133 genes are significantly differentially expressed (p < 0.05). Please note that 134 entries fall below the significance threshold as two entries correspond to different NCBI gene IDs, but the same En- sembl ID and were therefore treated as one gene.

Supplementary Table S3: List of significantly overrepresented KEGG pathways in the liver within the differentially expressed genes (DEGs). The *geneID* column contains the DEGs associated with each overrepresented KEGG pathway.

Supplementary Table S4: List of significantly overrepresented Gene Ontology (GO) terms in the liver within the differentially expressed genes (DEGs). The *geneID* column contains the DEGs associated with each overrepresented GO terms.

Supplementary Table S5: List of significantly enriched KEGG pathways in the liver according to genes ranked based on their p-value and the sign of the log2 fold-changes. The *core_enrichment* column contains the leading-edge genes, defined as the subset of genes that contributes most to the enrichment signal and drives the enrichment score within each enriched KEGG pathway’s gene set.

Supplementary Table S6: List of significantly enriched GO terms in the liver according to genes ranked based on their p-value and the sign of the log2 fold-changes. The *core_enrichment* column contains the leading-edge genes, defined as the subset of genes that contributes most to the enrichment signal and drives the enrichment score within each enriched GO terms gene set.

Supplementary Table S7: List of significantly overrepresented KEGG pathways in skeletal muscle within the differentially expressed genes (DEGs). The *geneID* column contains the DEGs associated with each overrepresented KEGG pathway.

Supplementary Table S8: List of significantly enriched KEGG pathways in skeletal muscle according to genes ranked based on their p-value and the sign of the log2 fold-changes. The *core_enrichment* column contains the leading-edge genes, defined as the subset of genes that contributes most to the enrichment signal and drives the enrichment score within each enriched KEGG pathway’s gene set.

Supplementary Table S9: List of significantly enriched GO terms in skeletal muscle according to genes ranked based on their p-value and the sign of the log2 fold-changes. The *core_enrichment* column contains the leading-edge genes, defined as the subset of genes that contributes most to the enrichment signal and drives the enrichment score within each enriched GO terms gene set.

## Appendix B. Supplementary Figures

Supplementary figures are available online (https://doi.org/10.5281/zenodo.20527211).

Supplementary Figure S1: Schematic overview of the iterative tree-based model selection procedure used to identify the simplest acceptable DESeq2 model.

Supplementary Figure S2: Principal component analysis and correlation plot of the read counts of all the liver samples (a—b) and skeletal muscle (c—d) included in this study, before any filtering was performed.

Supplementary Figure S3: Visualisation of enriched KEGG pathways in the liver. Genes are coloured according to their signed p-values. Genes in white were not detected in the RNA sequencing.

Supplementary Figure S4: Gene-concept network visualisation of GSEA results of the KEGG pathways in the liver. Orange nodes are pathways, and red and blue nodes are genes. Only genes common to more than one pathway are labelled.

Supplementary Figure S5: Gene-concept network visualisation of GSEA results of the Gene Ontology terms in the liver. Orange nodes are pathways, and red and blue nodes are genes. Only genes common to more than one pathway are labelled.

Supplementary Figure S6: Heatmap displaying the differentially expressed genes in muscle found in every over-represented KEGG pathways. Genes in blue are down-regulated in high-NUE pigs compared to low-NUE pigs while genes in red are up-regulated. Details on the genes are listed in Supplementary Table S7

Supplementary Figure S7: Visualisation of enriched KEGG pathways in skeletal muscle. Genes are coloured according to their signed p-values. Genes in white were not detected in the RNA sequencing.

Supplementary Figure S8: Gene-concept network visualisation of GSEA results of the KEGG pathways in skeletal muscle. Orange nodes are pathways and red and blue node are genes. Only genes common to more than 1 pathway are labeled

Supplementary Figure S9: Gene-concept network visualisation of GSEA results of the Gene Ontology terms in skeletal muscle. Orange nodes are pathways and red and blue node are genes. Only genes common to more than 1 pathway are labeled

## Acknowledgements

We thank Paolo Silacci’s team at Agroscope’s Animal Biology group for the extraction of RNA. We thank the Next Generation Sequencing Platform of the University of Bern for performing the high-throughput sequencing experiments, especially Pamela Nicholson and her team, and Marco Kreuzer, who provided valuable guidance with demultiplexing and the creation of the read count matrix. Gabriela Rudd-Garcés provided valuable comments on previous versions of this manuscript. Artificial intelligence tools were used to revise style and grammar of text (DeepL and ChatGPT), to find relevant literature (SciSpace) during the preparation of the manuscript and for the code to create figures. After using this tool, the authors reviewed and edited the content and take full responsibility for the content of the publication.

## Data, script, code, and supplementary information availability

Supplementary information, scripts and processed data required to reproduce the analyses are available online (https://doi.org/10.5281/zenodo.21776588). The raw RNA sequencing data have been deposited in the European Nucleotide Archive (ENA) under project accession PRJEB95768 (https://www.ebi.ac.uk/ena/browser/view/PRJEB95768).

